# Species responses to nutrient loading promote resistance but not temporal stability in floating macrophyte communities

**DOI:** 10.64898/2026.06.10.731482

**Authors:** Samuel R.P-J. Ross, Alexandru Mihai, Chii Kojima, David W. Armitage

**Author notes:** Correspondence: Samuel R.P-J. Ross, Graduate School of Agriculture, Kyoto University, Kitashirakawa Oiwake-cho, Sakyo-ku, Kyoto, 606-8502 Japan. E.

## Abstract

Determining the drivers of ecological stability amid accelerating global environmental change is a critical goal of contemporary ecology. Various candidate drivers have been suggested, with recent attention turning to response diversity—the variation among organism-environment responses. However, despite conceptual interest in response diversity as a driver of stability, there remain few field tests of this relationship. Using multi-species competitive communities of floating aquatic macrophytes as an experimental model for measuring temporal stability and response diversity to nutrient loading, we show that response diversity does not promote temporal stability of total macrophyte cover, but that communities with an uneven distribution of species responses were more resistant to an exogenous shock. To quantify macrophyte composition and growth dynamics from photographic time series of our experimental communities, we developed an open-source, scalable, machine learning workflow (*LeafMosaic*) capable of classifying four species from noisy field data including variable lighting, resolution, and plant morphology. We measured response diversity as the balance of positive and negative biomass growth responses to dissolved nitrate concentration, weighted by species’ relative contributions to biomass, and tested its effect on temporal stability and resistance to an unexpected pulse disturbance (a large typhoon that disrupted our outdoor mesocosms). Response imbalance predicted typhoon resistance, but species asynchrony and mean population stability best predicted community stability, with no direct or indirect effect of species responses. Overall, our results provide new experimental evidence for how the structure of species responses promotes stability, and we aim our *LeafMosaic* workflow to empower future field experiments using floating macrophytes to study response diversity and ecological stability.

## Introduction

The intensifying environmental pressures of the Anthropocene demand a mechanistic understanding of how ecosystems to respond to change. Ecological stability offers a multidimensional framework for understanding these responses (Donohue *et al*. 2013; Hillebrand *et al*. 2018), encompassing both community-level temporal stability and the system’s capacity to withstand and recover from discrete disturbances, as captured by metrics such as resistance and resilience. In natural ecosystems, more species-rich communities tend to have reduced total biomass variability—and therefore higher stability—through time (Yachi & Loreau 1999). Yet species richness alone does not confer this stability. Instead, a range of related mechanisms have been identified as important, including species interactions (Mougi & Kondoh 2012; Ratzke *et al*. 2020), asynchronous or compensatory dynamics (Morin *et al*. 2014; Shoemaker *et al*. 2022), and organismal traits (De Bello *et al*. 2021; Sasaki et al. 2026). Response diversity, an emergent property describing variation among organisms in their responses to environmental change (Elmqvist *et al*. 2003; Mori *et al*. 2013; Ross *et al*. 2023), has also been proposed as a key driver of ecological stability (Danet *et al*. 2024; Mori *et al*. 2013; Ross *et al*. 2026; Sasaki *et al*. 2019). Despite long-standing conceptual interest in response diversity as a mechanism promoting stability (Elmqvist *et al*. 2003; Nyström 2006), experimental tests of the response diversity-stability relationship remain rare, especially under field conditions (Ross & Sasaki 2024). Empirical evidence linking species’ environmental responses to community-level stability outcomes is therefore much needed.

Free-floating aquatic macrophytes are an effective experimental model for controlled laboratory experiments and semi-controlled field experiments (Dickinson & Miller 1998; Armitage & Jones 2019; Monacelli & Wilcox 2021; Couture et al. 2025). These plants are widespread, easy to manipulate, exhibit a range of life histories, and have short generation times, propagating asexually (Lemon et al. 2001; Armitage & Jones 2020; Jewell et al. 2023b). Floating macrophytes are also ecologically significant, often covering large extents of standing waters in lakes, ponds, and agricultural ditches (Duarte et al. 1986; Egertson et al. 2004), and have become noxious invasive species in many parts of the world. They also have applied uses in aquaculture, livestock feed, carbon capture, and bioremediation. Some species (*e.g.*, *Azolla* spp.) also fix nitrogen and contribute to the aquatic N cycle (Brouwer *et al*. 2017; Jewell et al. 2023a; Armitage et al. 2025; Kinerson et al. 2025). The diverse life history strategies and growth responses of floating aquatic macrophytes to environmental stressors, nutrients, and other factors (Henry-Silva et al. 2008; Silva et al. 2013; Si et al. 2019; Lanthemann & van Moorsel 2022) makes them ideal candidates for the study of response diversity and stability (McCann 2016). For example, models of aquatic macrophytes have demonstrated how variation in a response trait—shading tolerance—can affect the point beyond which a shallow lake transitions to a eutrophic state (Dakos et al. 2019). Under high levels of response diversity, the tipping point can even disappear, with the lake no longer exhibiting bistability. In a field experiment, submerged macrophyte communities with higher species diversity were more stable through time and better resisted algal bloom impacts on total biomass (Wang et al. 2025). However, this relationship was partly governed by a sampling effect, where dominant species with conservative nutrient use strategies could better regulate their elemental stoichiometry and resist exogenous nutrient disturbances. Floating aquatic macrophytes are known to exhibit interspecific differences in their responses to nutrient stoichiometry and temperature, knowledge of which can then be used to inform management when their overgrowth results in undesirable ecological regimes (McCann 2016).

Here we present results from an outdoor mesocosm experiment on four-species competitive communities of floating aquatic macrophytes in Okinawa, Japan, to empirically test the link between response diversity and community stability. To generate time series suitable for analysis we photographed mesocosm communities twice weekly and developed a machine learning workflow to classify plant species to estimate community composition. We present this open-source workflow here, and intend it to be useful for future studies of floating macrophytes (Mihai 2025). We predicted that higher community response diversity will reduce the temporal variability of total macrophyte biomass, because declines of sensitive species can be offset by neutral or positive responses of others (Yachi & Loreau 1999). Specifically, we measured species growth responses to nutrient loading in the form of nitrate concentration, which has a critical role in aquatic ecosystems; nitrate leaching is a key global change driver of eutrophication, water turbidity, plant die offs, and disruptions to aquatic food webs (Rabalais 2002). We quantify response diversity as the degree to which nitrate responses represent a balance of positive and negative responses in the community (Polazzo *et al*. 2025). This metric combines the direction and magnitude of species responses, where “imbalanced” communities are dominated by responses of a given direction (*e.g.*, almost all species respond negatively to nitrate addition), while “balanced” communities exhibit strong positive responses which offset equally strong negative responses (via asynchronous species dynamics), or because species have weak nitrate responses overall (and hence high population stability). By measuring the response diversity, species asynchrony, population stability, and community stability of macrophyte communities in a simple experimental test of the response diversity-stability relationship, we provide empirical evidence linking the structure of species’ environmental responses to community-level stability outcomes.

## Methods

### Floating macrophyte communities

We collected four species of floating macrophytes from agricultural ditches around sugarcane farmland in Ōyama and Kin on Okinawa main island, Japan (Figure 1). Our focal macrophyte species were the duckweeds *Lemna minor* (コウキクサ in Japanese) and *Spirodela polyrhiza* (ウキクサ) (Araceae), and the aquatic ferns *Salvinia molesta* (サンショウモ) and *Azolla cristata* (アカウキクサ) (Salviniaceae). These species encompass different sizes and functional groups of aquatic macrophytes, from the diminutive *L. minor* to the large *S. molesta*. The fern *A. cristata* is a nitrogen-fixing species, and, alongside *S. polyrhiza*, can produce dormant life stages in the forms of sporocarps and turions, respectively, which are mechanisms by which they can avoid stressful conditions. We chose this combination of species because they were *a priori* expected to have different tolerances to nutrient stress (McCann 2016), they are visually separable in multispecies communities (Figure 1), and regularly co-occur in the wild. These species are also primarily clonal and sourced from a single population, allowing us to reduce the intraspecific noise associated with species’ environmental responses.

**Figure 1.**
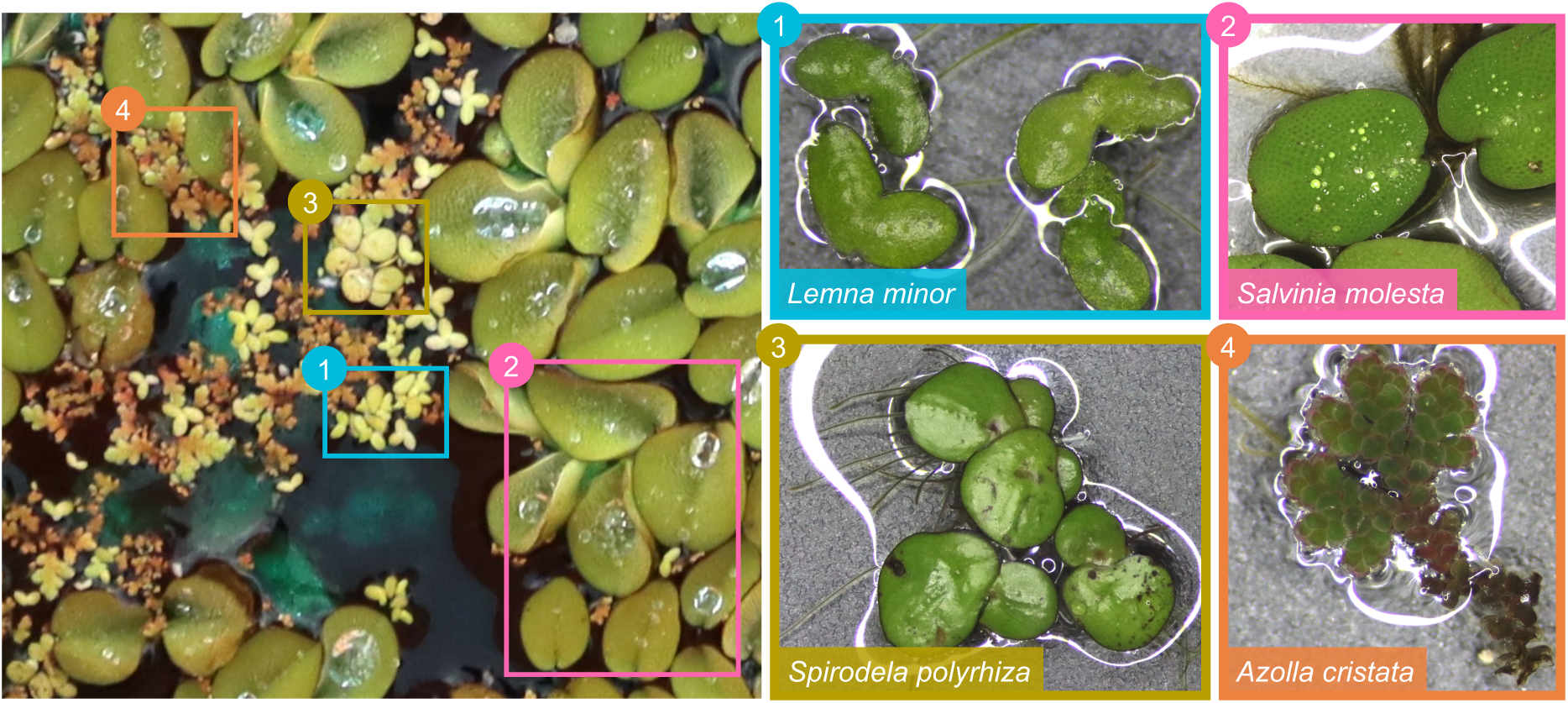
Left: a representative multi-species mesocosm community, with visually separable species. Right: the four species of floating aquatic macrophytes used in our experiment.

### Experimental design and timeline

Following washing under deionized water, macrophyte communities were established on 20 June 2023 (Day 0) and allowed to grow until 28 August 2023 (Day 70). Communities were housed in 100L polypropylene buckets (hereafter mesocosms; Figure 2a) filled with air temperature tap water to simulate realistic lentic bodies in Okinawa such as agricultural ponds and ditches. Mesocosms were established on an open gravel lot and were covered with a blue nylon mesh to exclude herbivorous pests. The mesh reduced light penetration by 4351 lx (± 2977 lx). Normal water temperatures throughout the 70-day study period were 31.0 °C (mean ±3.0 °C), ranging from 25.5 °C to 38.9 °C (Supplementary Figure S1). There was no temporal trend in temperature during the June-August Summer period in Okinawa, but a passing typhoon reduced water temperatures and temperature variability for ∼1 week (27.2 ±2.4 °C; see below). We added 0.1x concentrated Hoagland H-40 media without nitrogen to each mesocosm on Day 0 to facilitate initial plant growth (Supplementary Table S1). We chose this medium since it did not contain nitrogen, allowing us to independently manipulate nitrogen concentrations without affecting any other nutrients (Costa *et al*. 2009). Our study period coincided with the passing of Typhoon Khanun (Japan Meteorological Agency [JMA] 2023), a category 4 storm impacting Okinawa around 31 July (Day 41) to 6 August (Day 47). The typhoon reduced air and water temperatures (Figure S1) and resulted in unexpected and species-specific macrophyte mortality in many buckets (see *Results*).

**Figure 2.**
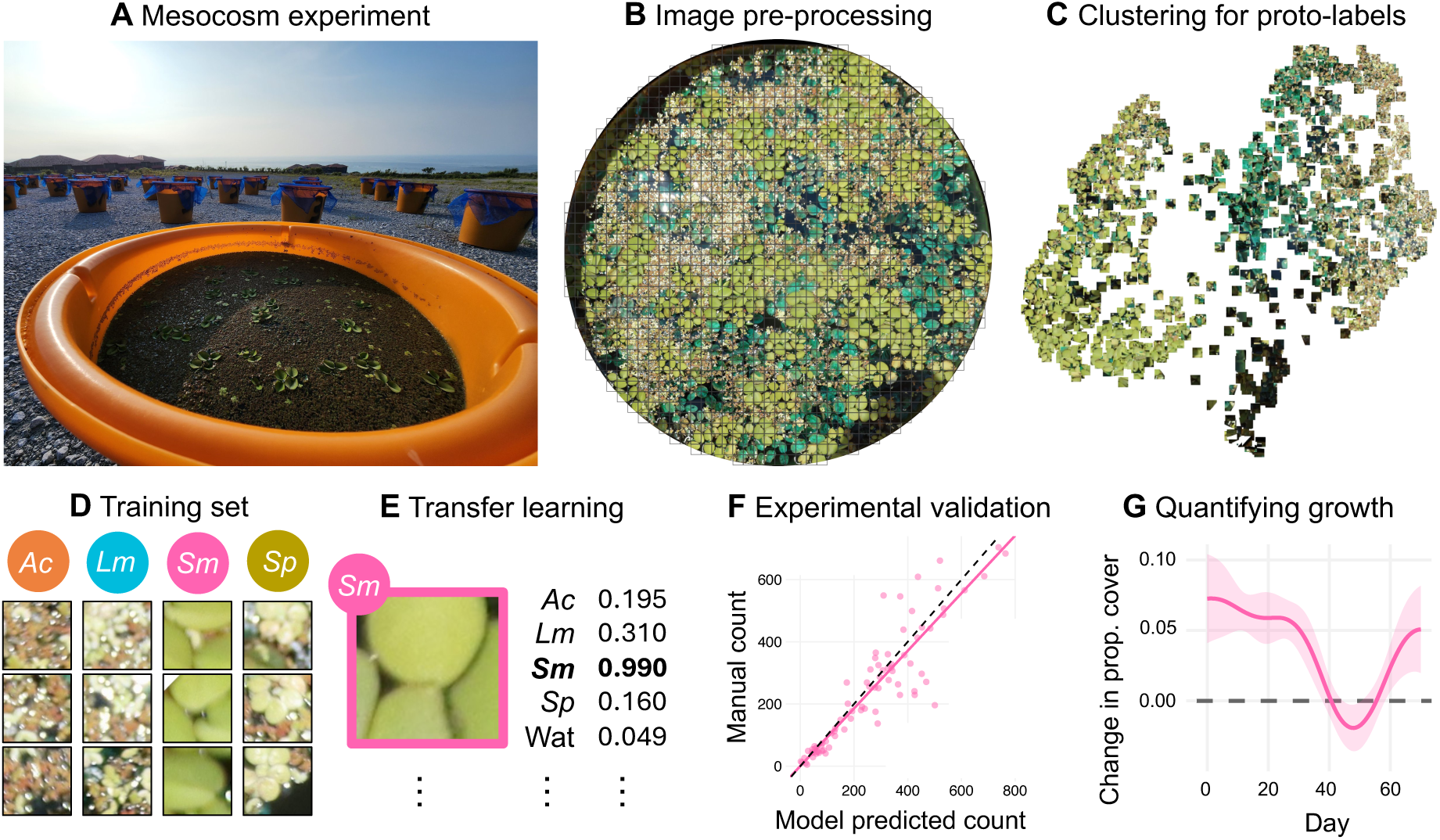
Workflow for image-based macrophyte classification with *LeafMosaic* (Mihai 2025). (A) Photograph of mesocosms. (B) Top-down photograph of the water surface, cropped, showing 100×100 pixel image tiles. (C) UMAP projection and HDBSCAN clustering of extracted features to generate proto-labels for training set construction. (D) Manual verification of clusters and assembly of species-specific training set. (E) Transfer learning classification using DenseNet and VGG models with fine-tuned classifier heads. (F) Experimental validation of transfer learning using manually classified data. (G) Quantification of species growth rates from time series of change in proportional cover data.

Since we measure response diversity as the diversity of growth responses to nutrient manipulation, we used a multifactorial experimental design. We crossed community composition (all combinations of 1-4 species richness levels) with nitrate (NO_3_^−^) enrichment (3 levels). We randomly assigned community composition and NO_3_^−^ concentration treatments to each mesocosm using a random number generator. Each macrophyte species was represented at each richness level six times for *S* = 1 (2 / NO_3_^−^ level), 6 times for *S* = 2 (2 / NO_3_^−^ level), 9 times for *S* = 3 (3 / NO_3_^−^ level), and 15 times for *S* = 4 (5 / NO_3_^−^ level). This design ensured we could derive growth rates for each species in isolation and in mixture under different treatments, allowing us to control for stochastic competitive exclusion scenarios in some mesocosms.

We exposed macrophyte communities to different nitrate (NO_3_^−^) concentrations by adding crystalline NaNO_3_ to mesocosms. Treatment levels were derived from empirical NO_3_^−^water samples from agricultural ponds and ditches around Okinawa where we sourced macrophytes. We chose three NO_3_^−^ treatment levels (low, medium, and high NO_3_^−^ addition) that represent realistic local NO_3_^−^ conditions in Okinawa while also differing sufficiently for treatment effects (Figure S2). To achieve and approximately maintain target NO_3_^−^concentrations for each treatment level, we pulsed NaNO_3_ additions every 7 days (9 pulses beginning Day 0). We added crystalline NaNO_3_ as appropriate for 100 L water volume to achieve a final concentration of ∼182, ∼656, and ∼1460 μg NO_3_^−^ L^−1^ for the low, medium, and high concentration treatments respectively (see Table S1). We stirred NaNO_3_ through the water column to facilitate even dissolving while minimising physical disturbance to the water surface.

To prevent macroalgal blooms, which can reduce nutrient availability for macrophyte growth, we added the commercially available “Aquashade” blue dye (Applied Biochemists, Germantown, USA) to each mesocosm. Aquashade dyes water blue and reduces phytoplankton growth in the water column by reducing photosynthetically available light in wavelengths of ∼600-650 nm (Batt *et al*. 2015; Tucker & Mischke 2020). On Day 0 we added 0.02 mL L^−1^ (20 ppm) to each mesocosm to dye water dark blue. We exceeded the manufacturer recommended 2.0 ppm Aquashade concentration because, unlike in ponds and lakes to which Aquashade is usually added, our mesocosms were exposed above ground to sunlight without shade, and mesocosms were light in colour, allowing light penetration via ground reflectance (Figure 2A). We topped up water levels *ad hoc* when they fell below 100 L due to evaporation and added 0.2 μL L^−1^ (0.2 ppm) to maintain Aquashade levels (Batt *et al*. 2015). Aquashade aids in suppressing macroalgal growth in the water column, but does not impact the growth of macrophytes on the water surface (Anderson & Martin 2005). Moreover, in support of previous work finding no effect on nitrite or total ammonia concentrations (Tucker & Mischke 2020), Aquashade did not alter nitrate availability or spectrophotometric detection ability in laboratory nitrogen assays here (polynomial regression coefficients ±95 confidence intervals overlapped for all tested Aquashade concentrations; Figure S3).

### Image segmentation and processing workflow using LeafMosaic

To assess macrophyte community change, we used a handheld Canon EOS 90D camera with EFS 18-55 mm lens (Canon Inc., Tokyo, Japan) to photograph the complete water surface of each mesocosm every 3-4 days; twice weekly photographs captured short-term change in macrophyte cover while balancing research effort in challenging field conditions. Including a gap in photography between 31 July and 8 August for Typhoon Khanun, during which fieldwork was not possible, our macrophyte community time series consisted of 20 photographs per bucket (1,379 total, 1 unusable data point) spanning 70 days (Figure S4).

Since accurately delineating and counting floating macrophytes from field photographs is very time-consuming and subjective among counters, we developed and applied a novel machine learning workflow to automatically identify and classify different macrophyte species and estimate macrophyte cover dynamics. Existing phenotyping workflows for floating macrophytes typically require highly controlled light and camera settings and aim at identifying individual plants in isolation (Kose et al. 2023). However, our images present a substantial challenge for automated machine learning models because this image set contains blurry photographs, those taken at inconsistent angles or directions, and in different light conditions (due to weather, photographer shadow, and time-of-day). Moreover, our macrophyte communities exhibited considerable phenotypic plasticity (particularly the colour and shape of *Salvinia molesta* and *Azolla cristata* (see Figure S4).

Our workflow contained the following steps (Figure 2), implemented in the newly developed *LeafMosaic* pipeline (Mihai 2025). To preprocess images for analysis, we first removed the background using the Rembg package (ver. 2.0.59; Gatis 2025) in Python (ver. 3.8; Python Software Foundation 2019), and manually fitted a circular crop to each image. We then split each photograph into 100×100-pixel square tiles (hereafter, *image tiles*; Figure 2B) and filtered out those with >50% background pixels. Tile-based approaches avoid the need for full-image segmentation and have been applied to estimate species cover from photographs (Körschens et al., 2021) and from UAV imagery of forest canopies (Nepi et al. 2025). Classification results were qualitatively equivalent regardless of the location of image tile delineation (Table S2).

To construct a training set, we first extracted features from each image tile using pre-trained DenseNet121 (Huang et al., 2017) and VGG16 (Simonyan & Zisserman, 2015) convolutional neural network (CNN) backbones with ImageNet weights (Russakovsky et al., 2015). Features derived from CNNs pre-trained on large general-purpose datasets transfer effectively to unsupervised classification tasks, often outperforming hand-engineered features (Guérin et al., 2018). We projected these high-dimensional feature vectors into two dimensions using UMAP (McInnes et al., 2018; Figure 2C) and applied HDBSCAN to the resulting embeddings to identify visually coherent clusters of tiles (Campello et al., 2013; McInnes et al., 2017). This combination of UMAP for dimensionality reduction and HDBSCAN for density-based clustering is effective for discovering structure in noisy, heterogeneous ecological data (Violet et al., 2025). In our workflow, unsupervised clustering served as a proto-labelling step, rapidly grouping tiles by visual similarity without the need for manual annotation. We did this for 40 photographs comprising a range of species combinations, phenotypes, and dates, and one author (S. Ross) reassigned incorrect species classifications as needed, ensuring species classifications always represented >50% cover of an image tile. Given the wide phenotypic variation in both *S. molesta* and *A. cristata*, we defined separate training classes for two phenotypes per species: green versus brown *S. molesta*, and green versus red *A. cristata*, plus classes for *L. minor, S. polyrhiza*, and non-target pixels (mostly water with <50% macrophyte cover), yielding eight classes total. Across 40 photographs, this produced a verified training set of 19,754 image tiles (Figure 2D): 7,393 *A. cristata*, 4,874 *S. molesta*, 2,854 *L. minor*, 2,560 *S. polyrhiza*, and 2,073 water tiles. See Supporting Methods and Table S3 for full details of the training dataset composition.

We next used the labeled training set for transfer learning, fine-tuning classifier heads on two architectures: DenseNet121 and VGG16. We retained fixed ImageNet feature extraction weights for DenseNet and VGG (Figure 2E). Transfer learning from ImageNet pretrained CNNs is standard for plant species classification, with DenseNet and VGG architectures consistently achieving high accuracy even on small, domain-specific training sets (Lee et al., 2023; Arun & Viknesh, 2025). Initial checks showed that models trained on all species at once had high misclassification rates for photographs with fewer than four species, while those trained on an individual species did not generalise well to multispecies communities. Accordingly, we trained a separate model for each unique 1-4 species combination of our focal species (15 combinations), and for each of our two architectures (see Supporting Methods, Table S4 and S5 for details).

We aimed to produce classifiers that could accurately estimate the relative abundance of different macrophyte species as well as their temporal dynamics—critical for accurately measuring stability. To validate our models, one author (S. Ross) manually reviewed a subset of classified image tiles (204,480 tiles from 160 photographs of 8 mesocosms), and manually assigned image tiles to species classifications, applying the same >50% cover threshold used when forming the training set. Accordingly, manual classifications produced more conservative abundance estimates than model classifications in most cases.

Finally, we assessed model performance of a range of models by fitting a linear regression to the manually verified (observed) tile counts per species per mesocosm per time point, and the model predicted counts (Figure 2F). For this experimental validation, models with 0.00 intercept, 1.00 slope, and high variance explained (adjusted *R^2^*) indicate that observed data are well predicted by the transfer learning model. We tested four combinations of model specifications and selected final model parameters based on these validation scores. 1) We compared the performance of models generated via transfer learning of DenseNet, VGG, and an arithmetic mean of the tile counts assigned to each species for a photo (model average). Ensemble approaches combining multiple CNN architectures can improve classification accuracy from individual models for plant identification tasks (Lee et al., 2023). 2) Tile counts were generated by models independently of time series position, missing the temporal autocorrelation of natural growth series. We therefore tested the effect of smoothing tile count time series using loess smoothers with alpha = 0.3, and compared results to raw machine learning outputs. 3) For each tile, we also estimated proportional macrophyte cover using HSV and LAB colour space filtering. We combined masks for green, yellow, and brown hue ranges with a lightness threshold and morphological noise reduction, and used these cover estimates alongside model predictions to filter out cases with <0.3 and <0.5 proportional cover within each image tile (see Supporting Methods and Table S5). 4) Finally, to fix the high-probability confusion of the visually similar *L. minor* and *S. polyrhiza* we tested whether filtering cases where *L. minor* and *S. polyrhiza* have classification confidence scores within a small window (<0.1 and <0.05) can improve performance. We filtered these cases by either removing the tile from the count if its probability confusion was within this threshold (*i.e*. reassigning the tile to the water class), or by randomly reassigning the tile to either *L. minor* or *S. polyrhiza* with equal probability.

### Measuring growth potential

To quantify response diversity to nitrate addition, we first needed to estimate species-specific growth under different NO_3_^−^ levels. We measured growth potential from generalised additive models (GAMs) of the natural logarithm of proportional macrophyte cover per species modelled against time (day since start), using the R package *mgcv* (ver 1.9-3; Wood 2011). Proportional cover is the proportion of 100×100 pixel tiles classified as a given species per mesocosm per day (out of the total number of tiles per mesocosm per day). We clipped time series to only include data from time points before total macrophyte cover (all species) reached 90% of the water surface, thereby avoiding measuring growth when with limited by competition (Figure 2G). Data were best described by a beta distribution, so we modelled GAMs with the beta family and logit link function. We modelled time as a Gaussian process smooth with 8 basis functions (*k*), since *k* = 8 had the lowest Akaike information criterion (corrected for sample size) across all mesocosms (ΔAICc = 3.87; Figure S5).

We calculated the first derivative of each smoothed time series, yielding a series of changes in log proportional cover per species per mesocosm through time (Figure S6). From these time series, we measured average growth rate as the arithmetic mean of the derivatives of the log proportional cover time series per species per mesocosm (Shipley & Hunt 1996). We then standardised these growth rates by each species’ maximum proportional cover per mesocosm to ensure growth potential values were ecologically meaningful (Figure 3). The resulting growth potential values for each species and NO_3_^−^treatment combination qualitatively matched previous work and the natural history of floating aquatic macrophytes (Armitage & Jones 2019; Lemon et al. 2001; Vermaat & Hanif 1998), well reflected realised growth rates observed during the experiment (S. Ross, *pers. obs.*), and exhibited sufficient variation between species for quantifying response diversity (Figure 3).

**Figure 3.**
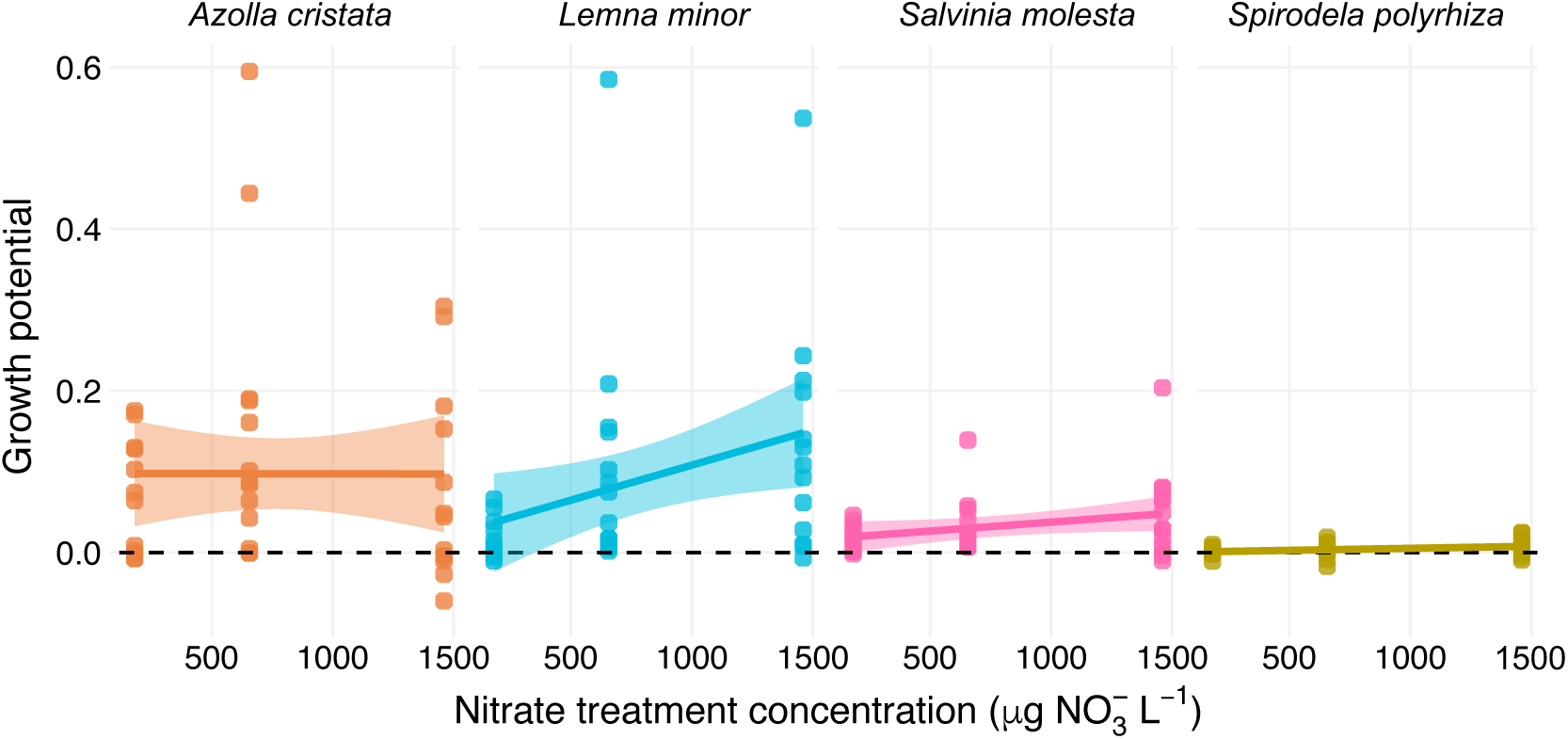
Variation in species growth potential at different nitrate (NO_3–_) concentration treatments. Species-specific growth potential measured as the arithmetic mean of derivatives of log proportional cover through time, divided by maximum within-mesocosm proportional cover. Data were measured under three pulsed nitrate NO_3_^−^ addition treatments: low (∼182 μg NO_3_^−^ L^−1^), medium (∼656 μg NO_3_^−^ L^−1^), and high concentration (∼1460 μg NO_3_^−^ L^−1^).

### Response diversity, population stability, and species asynchrony

From the multi-species proportional cover time series and derived growth potential values, we measured several biotic variables which we expected to mechanistically drive community stability: mean growth potential, response diversity, mean population stability, and species asynchrony. As mean species responses can predict stability and coexistence (de Laender et al. 2023; Kunze et al. 2026), we measured the mean growth potential for each mesocosm as the arithmetic mean of the growth potential values of all species in the community. By measuring this from the mesocosm-specific growth potential values, this metric captures the realised average growth response in each macrophyte community.

We used the response imbalance metric to measure response diversity (Polazzo et al. 2025). This metric unifies efforts to separately measure the magnitude versus the direction of organisms’ environmental response functions (Ross *et al*. 2023). Response diversity is typically considered along environmental gradients (*e.g.*, Cariveau *et al*. 2013), so here we used species’ growth responses along a nitrate enrichment gradient from low to high NO_3_^−^ concentration (Figure 3), to measure response imbalance per mesocosm as:

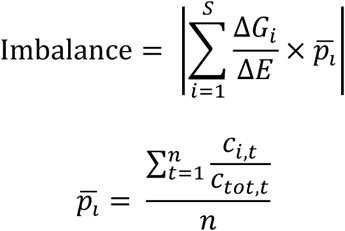

where: Δ*G*_*i*_ is the change in growth potential of species *i*; Δ*E* is the change in NO_3_^−^ concentration between the high and low NO_3_^−^ treatments (always 1278 μg NO_3_^−^ L^−1^ here); *S* is the species richness of the community; *p̅*_*ᶥ*_ is the average contribution to total macrophyte cover; *n* is the time series length (70 days); *c*_*i*,*t*_ is the proportional cover of species *i* on day *t*; and *c*_*tot*,*t*_ is the total proportional cover of all species combined on day *t*. By scaling imbalance by *p̅*_*ᶥ*_, we measured a realised version of response diversity which accounts for species-specific contributions to the total macrophyte community cover (Polazzo et al. 2025).

Response imbalance scales from zero to infinity, where higher values represent communities with more “imbalanced” NO_3_^−^ responses (that is, those dominated by either positive or negative responses to NO_3_^−^ concentration). On the other hand, low values of imbalance can arise in two ways: (1) strong positive NO_3_^−^ responses are counterbalanced by equally strong negative ones, resulting in stability via asynchronous species dynamics (Loreau & de Mazancourt 2013), or (2) responses to NO_3_^−^ enrichment are, on average, weak, resulting in high population stability and, in turn, community stability (Thibaut & Connolly 2013). In this way, low values of response imbalance (high “balance” or weak response) are expected to contribute to community stability; that is, we expect a negative relationship between response imbalance and community stability (Polazzo et al. 2025).

We measured species asynchrony (1 − *ϕ*; Loreau & de Mazancourt 2008) for each mesocosm with species richness >1 as 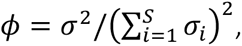 where *σ*^2^ is the variance of total macrophyte cover (across all species through time), and *σ*_*i*_is the temporal standard deviation of proportional cover for each species *i* of all S species. Asynchrony (1 − *ϕ*) then ranges from zero to one, representing assemblages with perfectly synchronised proportional cover fluctuations among species (0) to complete asynchrony (1; Sasaki et al. 2019).

To measure mean population stability, we use the inverse of Thibaut & Connolly’s (2013) average species-level coefficient of variation weighted by species’ relative proportional cover 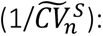

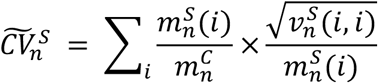

where: 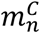 is the temporal mean of total macrophyte cover (*i.e.*, the sum of species-level mean cover, 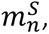 for all species); 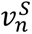 is the variance of macrophyte cover for species *i* such that 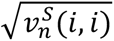 represents the temporal standard deviation of macrophyte cover. By using the inverse of 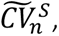 higher values of mean population stability represent cases where species vary less through time relative to their temporal means.

### Temporal community stability and typhoon resistance

For each mesocosm, we calculated temporal community stability using the untransformed total macrophyte cover across the whole experiment (20 time points spanning 70 days). We measured community stability of each macrophyte cover time series as the ratio of the temporal mean to the temporal standard deviation (µ/σ), the inverse of the coefficient of variation (Lehman & Tilman 2000; Isbell et al. 2009).

Our study period coincided with the landfall of category 4 supertyphoon Khanun (31 July–6 August, experiment days 41–47), the largest to hit Okinawa since 2018 (JMA 2023; Ross *et al*. 2024). Field observations suggested changes to relative macrophyte cover following the typhoon (S. Ross & C. Kojima, *pers. obs.*), offering a unique opportunity to explore the potential for response diversity to prime macrophyte communities against an exogenous pulse disturbance. We thus measured resistance to possible typhoon disturbance for total macrophyte cover and separately for each species as the instantaneous log response ratio (Hillebrand *et al*. 2018) between the disturbed state (an arithmetic mean across three dates: 8, 10, 15 August 2023) and the pre-disturbance state (25, 28, 31 July 2023).

### Data analysis

Data were analysed in the R programming language (ver. 4.5.1; R Core Team 2025), using packages *glmmTMB* (ver. 1.1.14; Brooks et al. 2017), *DHARMa* (ver. 0.4.7; Hartig 2024), and *piecewiseSEM* (ver. 2.3.1; Lefcheck 2016). Continuous predictors (mean growth potential, response imbalance, species asynchrony, and mean population stability) were z-transformed to standardised effect sizes using the formula 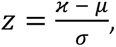 where *ϰ* is a value of a continuous variable and *μ* and *σ* are the mean and standard deviation of all values of that variable, respectively.

To test which predictor variables best explained stability outcomes, we first fitted a generalised linear mixed model (GLMM) with community stability as the response variable, and mean growth potential, response imbalance, species asynchrony, and mean population stability as predictor variables. NO_3_^−^ treatment (a factor with three levels: low, medium, or high NO_3_^−^) was included as an interaction term, allowing the slope and intercept of mean growth potential, response imbalance, species asynchrony, and mean population stability to vary by NO_3_^−^ treatment. We included species richness as a random effect to account for any direct effect of richness on stability. We used a model selection approach to choose between this full model, and models with subsets of variables, using the “dredge” function in the *MuMIn* R package (ver. 1.48.11; Bartoń 2025). Models were fitted using a Gaussian distribution with identity link function, and evaluated via the “simulateResiduals” function in *glmmTMB*. We used the small sample-corrected Akaike Information Criterion (AICc) to compare models, treating candidate models with ΔAICc ≤ 4 as having equivalent support. We used maximum likelihood estimation for model selection, followed by weighted model averaging across all candidate models with ΔAICc ≤ 4. Model-averaged parameter estimates and variable importance were weighted based on AICc-derived model weights. We used a conservative “full” rather than “conditional” model average, since conditional averages bias coefficients away from zero by only averaging over models that include a given term (Lukacs et al. 2009).

We further tested the direct and indirect pathways through which the above predictor variables can impact community stability using a piecewise structural equation model (SEM) framework (Lefcheck 2016). Paths included in the full piecewise SEM were informed by the ecological literature on response diversity and stability: realised response imbalance was hypothesised to have both direct and indirect effects on stability, via species asynchrony and mean population stability (Sasaki et al. 2019; White et al. 2023; Polazzo et al. 2025). We first expected mean growth potential to have only a direct effect on community stability, but tests of directed separation identified indirect pathways from mean growth potential through both species asynchrony and mean population stability as important. We thus included these additional paths in our SEM. The full SEM therefore comprised three models: community stability predicted by mean growth potential, realised response imbalance, species asynchrony, and mean population stability; species asynchrony predicted by realised response imbalance and mean growth potential; and mean population stability also predicted by realised response imbalance and mean growth potential (see Figure S7). All paths within the SEM were fitted using generalised linear models (GLMs) with Gaussian error structure and the identity link function, and global goodness-of-fit evaluated using Fisher’s C statistic with a threshold of *p* > 0.05 indicating a valid piecewise SEM model (Shipley, 2009). In all cases we used standardised estimates of coefficients to plot associations between variables. Given that piecewise SEM workflows cannot easily handle categorical data, we first constructed the above SEM on the entire dataset, and then on three data subsets representing only the mesocosms in the low, medium, and high NO_3_^−^ enrichment treatments. Qualitative differences in the structure of piecewise SEMs between the three NO_3_^−^ treatments would then suggest effects of NO_3_^−^ on direct and indirect stability drivers.

Finally, we tested the effect of realised response imbalance, mean growth potential, NO_3_^−^ enrichment treatment, and the interaction between imbalance and NO_3_^−^ treatment and mean growth potential and NO_3_^−^ treatment on community resistance (log response ratio [LRR] of total macrophyte cover). We did not use mean population stability or species asynchrony as predictors of resistance, since their role in resistance to discrete disturbance is less clear than in stabilising community dynamics through time (Gonzalez & Loreau 2009). We again fitted a full GLMM with Gaussian family and identity link function, including species richness as a random effect, then did model selection with AICc to identify the most parsimonious models based on ΔAICc ≤ 4. We then conducted weighted model averaging across candidate models with ΔAICc ≤ 4 using AICc-derived model weights and a “full” rather than “conditional” model average. Finally, to determine whether species differed in their resistance to typhoon disturbance, we used ANOVA with an interaction between species identity and NO_3_^−^ enrichment treatment as predictors and macrophyte resistance as the response variable. In this case, macrophyte resistance was measured as the per species instantaneous LRR on macrophyte cover and thus is equivalent to “population” rather than community resistance.

## Results

### Model classifier performance

Within the transfer learning training-testing framework, models returned a mean classification accuracy of 91.8% across all model-by-species combinations, with DenseNet121 performing better (92.5%) than VGG16 (90.9%; Table S2). Per-species accuracy ranged from 89.1% for *S. molesta* to 93.0% for *L. minor* (Table S3), with modest degradation as species richness increased from single-species classifiers (93.3%) to four-species mixtures (90.0%).

Our *LeafMosaic* workflow produced a total of 2,025,145 image tiles which were classified into our four focal species or background (water) classes. Experimental validation revealed that both *Azolla cristata* and *Salvinia molesta* were best classified by a model average between VGG and DenseNet, with a loess smoother on the time series, and image tiles removed if they contained <0.3 within-tile macrophyte cover (Table S7). The *S. molesta* model (slope [*m*] = 0.93, intercept [*b*] = 0.47, adjusted *R^2^* = 0.91; Figure S8C) performed marginally better than the *A. cristata* model (*m* = 0.89, *b* = 6.2, *R^2^* = 0.86; Figure S8A). The *Lemna minor* and *Spirodela polyrhiza* models with best classification accuracy were both DenseNet models with a loess smoother on the time series, and without any within-tile cover adjustments (Table S7). Both species were best classified when including a post-processing step to adjust cases with high confusion scores between *L. minor* and *S. polyrhiza*; the *L. minor* model randomly reassigned image tiles as *L. minor* or *S. polyrhiza* when their classification probability was within 0.05 for both species, while the best *S. polyrhiza* model instead removed those image tiles. The final *L. minor* model (*m* = 0.74, *b* = 18.4, *R^2^* = 0.84; Figure S8B) outperformed the *S. polyrhiza* model (*m* = 0.65, *b* = 1.85, *R^2^* = 0.85; Figure S8D). Critically, visual checks of time series confirmed that for all species, macrophyte cover dynamics well matched those from manual classification (Figure S9).

### Temporal community stability

Model selection on the model of community stability resulted in 9 candidate models with ΔAICc ≤ 4. The cumulative sum of model weights of these models was 0.87. Effects of species asynchrony and mean population stability were included in all 9 candidate models, mean growth potential was included in 6 candidate models, and realised response imbalance in 2. NO_3_^−^ treatment was included as a main effect in 5 candidate models, while its interactions with mean growth potential and mean population stability were included in 2 candidate models.

Our weighted model average of the nine candidate models included significant effects of species asynchrony and mean population on community stability with similar effect sizes (Figure 4A). The partial effect of species asynchrony on community stability was positive (Standardised effect size [*z*] = 0.41 ± 0.06 Adjusted Standard Error); communities with less synchronised species dynamics were more stable through time. The partial effect of mean population stability on community stability was also positive (*z* = 0.40 ± 0.07); communities with less strongly fluctuating species cover were more stable in aggregate cover through time. In contrast, our model average did not include a significant direct effect of mean growth potential (*z* = −0.04 ± 0.07) or realised response imbalance (*z* = 0.005 ± 0.02) on community stability (Figure 4A).

**Figure 4.**
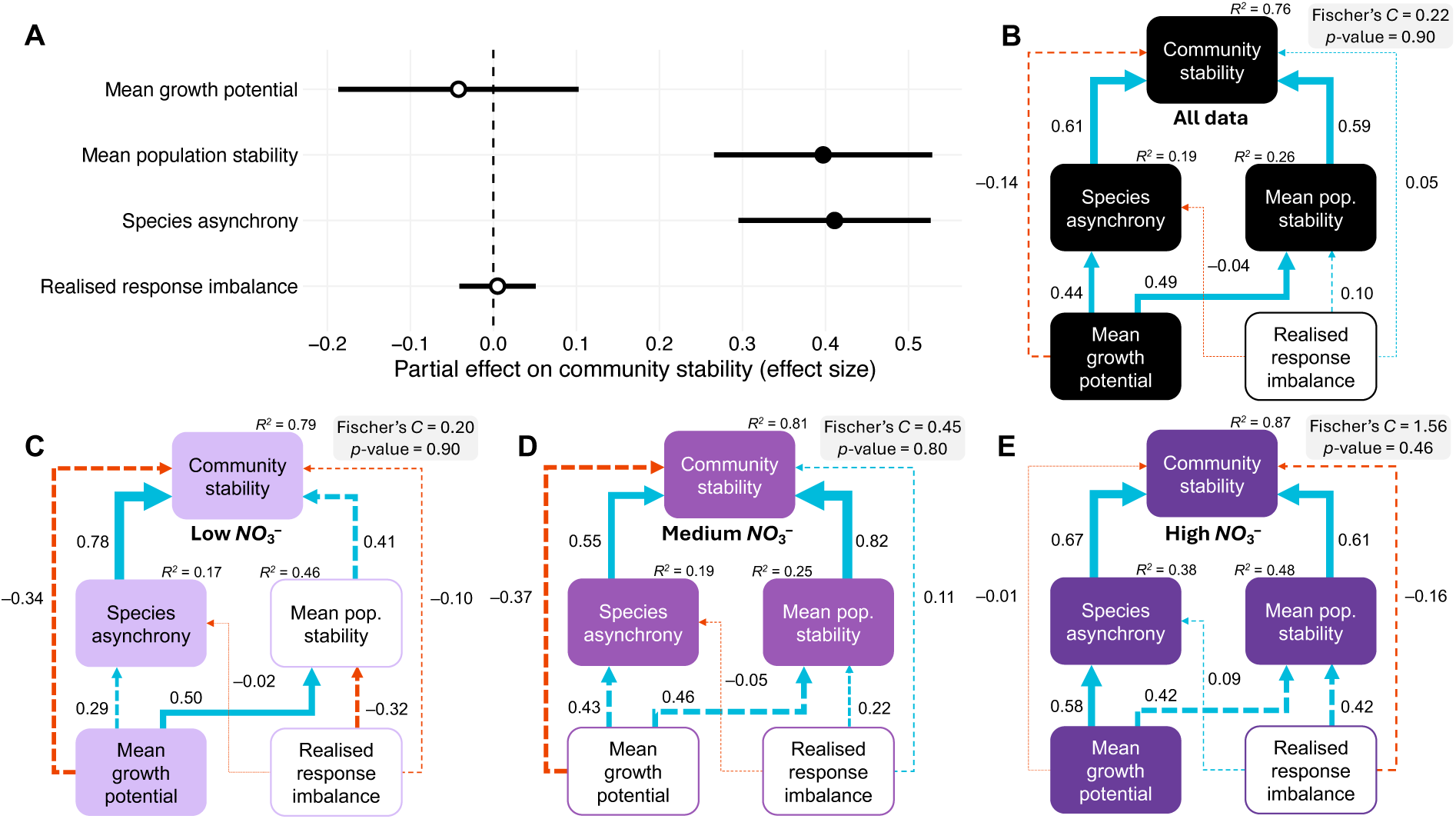
Direct and indirect predictors of temporal community stability in experimental macrophyte communities. A) Partial effects of four biotic predictors (mean growth potential, mean population stability, species asynchrony, and realised response imbalance) on community stability, measured as standardised effect sizes of the slope of the relationship in a weighted model average of nine closely competing candidate mixed effects models (see Methods). B) Piecewise structural equation model (SEM) on the whole dataset to identify possible indirect pathways driving community stability. We also built SEMs on subsets of the data to disentangle possible NO_3_^−^ treatment effects on the drivers of stability: C) the low NO_3_^−^ treatment (∼182 μg NO_3_^−^ L^−1^); D) the medium NO_3_^−^ treatment (∼656 μg NO_3_^−^ L^−1^); and E) the high NO_3_^−^ treatment (∼1460 μg NO_3_^−^ L^−1^). Solid arrows indicate significant correlations between pairs of variables, dashed arrows indicate nonsignificant relationships. Blue arrows and positive standardised estimates of coefficients indicate a positive relationship between two variables, while red arrows and negative values are negative relationships. Arrow width scales with the size of standardised model coefficients for visualisation. R^2^ values indicate the variance explained by all direct paths into a variable. We tested goodness-of-fit with Fischer’s tests, where nonsignificant values (*p* > 0.05) indicate suitable model fit. All paths included in full piecewise SEMs were informed by relevant literature (Figure S7).

Structural equation models (SEMs) to separate direct and indirect effects of predictor variables on community stability found consistent direct positive effects of both species asynchrony and mean population stability on community stability (Figure 4B,D,E). However, in the Low NO_3_^−^ treatment, mean population stability was not significantly related to community stability (Figure 4C). The SEM on the full dataset supported the notion that mean growth potential did not directly affect community stability, but was indirectly positively related to community stability through its positive relationship with species asynchrony and mean population stability (Figure 4B). This effect was less clear on the NO_3_^−^ treatment data subsets: mean growth potential was not directly or indirectly related to community stability in the low or medium NO_3_^−^ treatment communities, but was related indirectly to community stability at high NO_3_^−^ concentration, only through the species asynchrony pathway (Figure 4E). Contrary to our expectations, realised response imbalance was not directly or indirectly related to community stability in any of the models.

### Resistance to typhoon disturbance

Model selection on the model of community resistance (Log Response Ratio [LRR] of immediately before versus after the typhoon disturbance) resulted in 3 candidate models with ΔAICc ≤ 4. The cumulative sum of model weights of these models was 0.83. Realised response imbalance was included as a predictor in all 3 candidate models, mean growth potential was included in one model, and NO_3_^−^ treatment in one. NO_3_^−^ treatment interactions with other variables were not included in the candidate model set.

Our weighted model average of the three candidate models included no significant effect of mean growth potential (Standardised effect size [*z*] = −0.001 ± 0.03 Adjusted Standard Error; Figure 5A) or NO_3_^−^ treatment (ANOVA_2,42_: *F* = 0.62, *p* = 0.54; Figure S10) on community resistance, but a significant effect of realised response imbalance. The partial effect of response imbalance on community resistance was positive (*z* = 0.20 ± 0.08; Figure 5A), suggesting that communities with less balanced NO_3_^−^ responses were more resistant to typhoon disturbance, after accounting for the relative contribution of each species to total macrophyte community cover (Figure S11). However, this effect seemingly was not driven solely by species contributions to cover; there was no significant difference in realised response imbalance between communities with high versus low evenness in proportional cover (T-test: *t* = 0.84, *df* = 27.6, *p* = 0.41; Figure S12).

**Figure 5.**
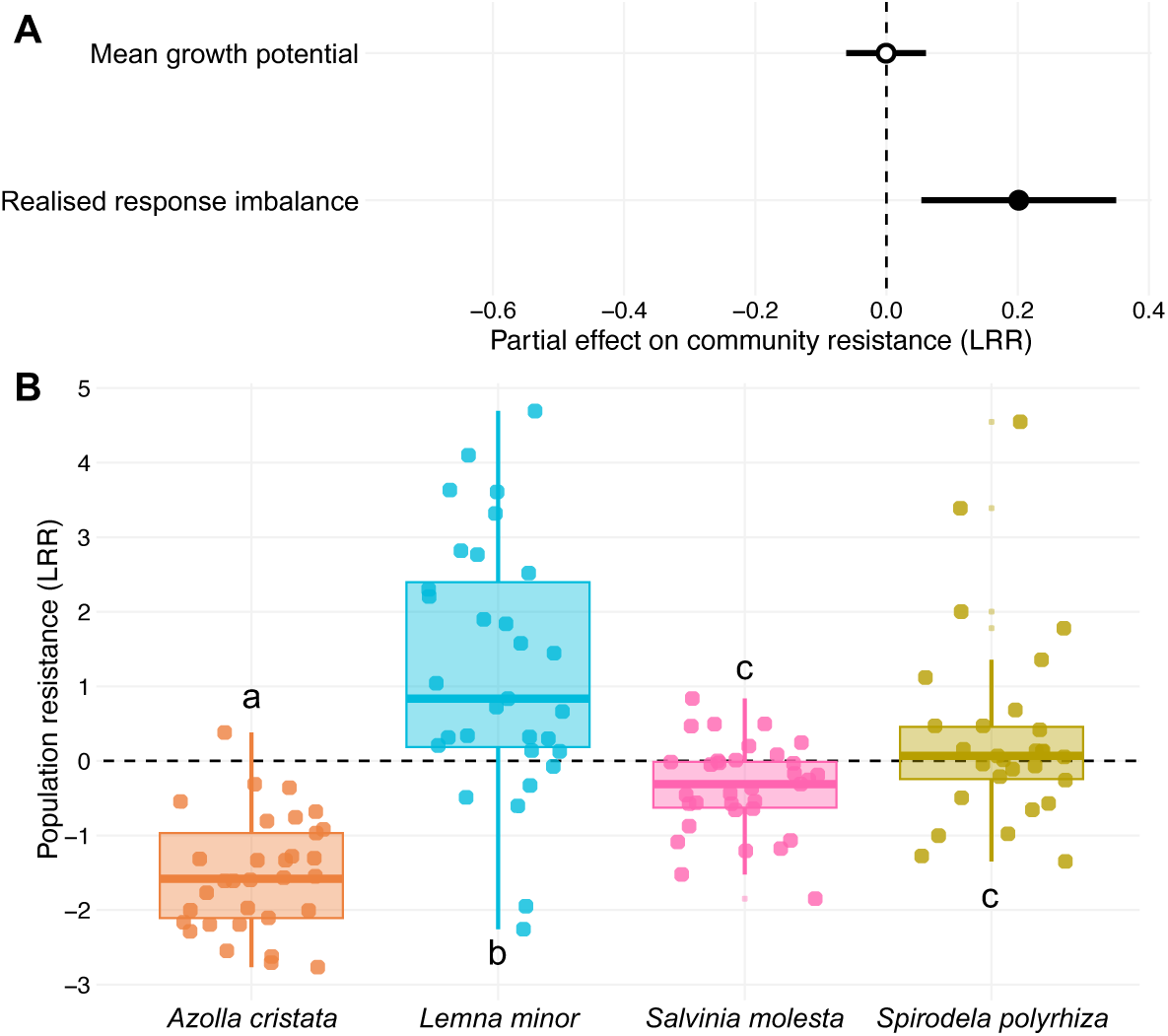
Predictors of community and population resistance in experimental macrophyte communities. A) Partial effects of mean growth potential and realised response imbalance on community resistance (log response ratio [LRR] of the change in total macrophyte community cover), measured as standardised effect sizes of the slope of the relationship in a weighted model average of three closely competing candidate mixed effects models (see Methods). B) Significant effect of species identity on population resistance, measured as the LRR of the change in species-specific cover following typhoon disturbance. Values above zero indicate a post-typhoon increase in macrophyte cover per species, while values below zero indicate a decrease. Lowercase letters denote significant pairwise contrasts from Tukey HSD tests.

We also measured population resistance as the species-specific change in macrophyte cover (LRR) following typhoon disturbance. As with community resistance, we found no effect of NO_3_^−^ treatment on macrophyte population resistance (ANOVA_2,113_: *F* = 1.72, *p* = 0.18; Figure S13). However, we found that species differed significantly in their species-level (population) change in macrophyte cover (ANOVA_3,113_: *F* = 31.1, *p* < 0.001). Post-hoc Tukey HSD tests revealed that all pairwise contrasts between our four macrophyte species were significant except for between *Salvinia molesta* and *Spirodela polyrhiza*, both of which responded quite weakly to typhoon disturbance (Figure 5B). The proportional macrophyte cover of *Azolla cristata* declined significantly following typhoon disturbance, while *Lemna minor* cover increased. There was no evidence for an interactive effect of NO_3_^−^ treatment and species identity on population resistance (ANOVA_6,113_: *F* = 0.45, *p* = 0.84).

## Discussion

Here we experimentally tested how the structure of species-environment responses affects the stability of competitive communities of floating aquatic macrophytes. Specifically we asked whether a balance of positive versus negative growth responses to nitrate (NO_3_^−^) enrichment can stabilise total macrophyte cover through time, and in response to an unexpected exogenous shock (a large typhoon). Despite conceptual interest in species-environment relationships and response diversity for years (Elmqvist *et al*. 2003; Ives *et al*. 1999; Ives & Carpenter 2007; Mori *et al*. 2013), there are still few field tests of the stabilising potential of response diversity (Ross & Sasaki 2024). Recent conceptual and methodological developments have sought to address this gap, operationalising the response diversity concept for empirical study (Ross et al. 2023; Polazzo et al. 2024, 2025). Leveraging these developments, we tested how species’ NO_3_^−^ responses shape stability outcomes in outdoor mesocosm communities, finding that species responses did not confer temporal stability but that the distribution of responses affected community resistance to a large pulse disturbance.

We found no effect of the balance of species responses to NO_3_^−^ enrichment on the temporal stability of total macrophyte cover. Responses to NO_3_^−^ can be balanced either because strong positive responses are offset by negative ones, resulting in species asynchrony, or because NO_3_^−^ responses are, on average, weak (Polazzo et al. 2025). Both these effects—species asynchrony and mean population stability—can be important drivers of temporal stability at the community level (Yachi & Loreau 1999; Thibaut & Connolly 2013; Sasaki et al. 2024; Wang et al. 2024). Accordingly, we tested whether species responses indirectly drove community stability, as expected by response diversity theory (Ross & Sasaki 2024; Polazzo et al. 2025). However, we found no indirect or direct effect of the balance of species’ NO_3_^−^ responses on community stability. Instead, we found that both species asynchrony and mean population stability were consistently positively related to community stability, and that mean species growth responses, rather than the distribution or balance of these responses, promoted the stabilising role of species asynchrony and mean population stability on total macrophyte cover. This result is consistent with the finding that species mean responses to temperature, rather than response diversity, drove community stability in model competitive communities (Kunze et al. 2026). Yet, in the absence of interspecific interactions, Kunze *et al*. (2026) found response diversity can be stabilising. Though species’ environmental responses may be more important for shaping community outcomes than species interactions (Houlahan et al. 2007; Ives & Carpenter 2007), perhaps competition between floating macrophytes reduced the potential for response diversity to confer community stability in our multispecies mesocosms.

Higher mean growth potential was related to both higher species asynchrony and higher mean population stability in our study. Faster growth rates can result in population or species asynchrony in spatially structured models (Heino et al. 1997; Matter 2001) and empirical communities (*e.g.*, Li et al. 2021). Mechanistically, this can occur through the Moran effect; in globally stochastic environments, populations fluctuate synchronously if environmental responses are identical, but high growth rates tend to chaotic dynamics, promoting asynchrony even in response to shared environmental drivers (see Heino et al. 1997). Fast growing species also tend to have high reactivity to environmental changes, allowing these populations to buffer environmental stochasticity and remain stable, in contrast to those in stable environments where slow-growing species are otherwise expected to have more stable populations (Pribil & Houlahan 2003). Given our outdoor mesocosm communities were exposed to natural daily temperature and precipitation regimes, this may in part explain why higher mean growth potential was related to higher mean population stability. Moreover, when smaller organisms have high reproductive potential—as is the case with *Lemna minor* here (see also Vermaat & Hanif 1998; McCann 2016)—their high carrying capacity per unit area reduces the strength of demographic stochasticity and can thus stabilise population dynamics (de Mazancourt et al. 2013). Taken together, our results support the idea that higher population growth rates promote community stability simultaneously through species asynchrony and mean population stability (Li et al. 2021).

Our focal macrophyte species had different biomass responses to an unexpected exogenous shock in the form of a large typhoon (JMA 2023). Typhoons alter water turbidity and the chemical composition of freshwaters through rainwater and runoff from surrounding soils and stream discharge (Peierls et al. 2003; Stockwell et al. 2020; Santiago-Vera & Ramírez 2026), and in Okinawa faster wind speeds during typhoons are associated with salination from seawater (Sakihama & Tokuyama 2005) and nutrient input from pre-senescent leaf fall (Xu et al. 2004). Our mesocosm communities were covered with a nylon mesh which prevented wind- or animal-assisted immigration or emigration and water level was maintained despite rainfall via an outflow hole smaller than any macrophyte fronds. Moreover, aboveground mesocosms were not subject to chemical runoff from soils, did not deliberately include trophic interactions with macroinvertebrates or other aquatic herbivores, and algal blooms were suppressed by macrophyte cover and use of Aquashade dye. Thus, typhoon effects on our experimental plants could only occur via physical disturbance and changes to water chemistry, which in turn can be accompanied by meiofaunal and bacterial blooms in the water column (Lai et al. 2021; Santiago-Vera & Ramírez 2026). In Hainan Island, China, typhoons altered the morphology of free-floating macrophytes; plants invest less in stem height and more in belowground traits to limit mechanical damage in typhoon-prone areas (Wang et al. 2010). In the same system, Wang *et al*. (2008) found that typhoon disturbance increases macrophyte richness through an increase in structural habitat heterogeneity, and niche differentiation, maintained by regular typhoon disturbance. In our system, the proportional cover of *Azolla cristata* declined after the typhoon, as did *Salvinia molesta*, which was submerged for several days before re-emerging (S. Ross, *pers. obs.*). Both these species have taller, more emergent growth forms relative to *Spirodela polyrhiza* and *Lemna minor*, with increased exposure to mechanical disturbance. In contrast, the flat leaves of *S. polyrhiza* may explain its lack of typhoon response, while *L. minor*, with its high growth potential can exploit the competitive release caused by biomass declines or submergence of its *A. cristata* and *S. molesta* competitors (Couture et al. 2025). Alternatively, phosphorous and nitrogen pulses caused by typhoons (Gao et al. 2021) may benefit resource-acquisitive macrophytes such as *Lemna minor*, which had higher growth potential under our high NO_3_^−^ enrichment treatment (McCann 2016; Wang *et al*. 2025). In lentic waterbodies, nutrient enrichment can shift ecosystems to an alternate state dominated by floating macrophytes (Scheffer et al. 2003). Differences between macrophyte species in their biomass growth responses to mechanical disturbance and nutrient input (Wang et al. 2010; McCann 2016) may therefore be exacerbated by typhoon disturbance, increasing the probability of transitions to macrophyte-dominated states in Okinawa’s freshwater systems subjected to frequent typhoons.

The structure of species’ NO_3_^−^ responses drove community resistance but not community stability. This supports the notion that response diversity may confer stability for dimensions beyond the temporal variability most often considered in response diversity studies (Mori et al. 2013; Ross et al. 2026). Theory predicts that a balance of positive and negative species responses to the environment should promote stability (Yachi & Loreau 1999; Polazzo et al. 2025). Yet, we found the opposite. Higher values of realised response imbalance—representing communities dominated by NO_3_^−^ responses of a given direction—were related to greater community resistance. Realised response imbalance considers the balance of species’ environmental responses in their realised ecological context by weighting species responses by their relative abundances (Polazzo et al. 2025). As such, dominant species with beneficial typhoon responses could explain this effect, such as the high growth potential and positive NO_3_^−^ response of the duckweed *L. minor*. Given their disproportionate contribution to community biomass, dominant species can be critical determinants of stability in a range of systems (Sasaki & Lauenroth 2011; Kunze et al. 2025; Xu et al. 2026). However, there was no effect of species evenness on the realised response imbalance of our experimental communities. Our results thus suggest that communities of species with imbalanced responses, but not dominant biomass, confer greater resistance to typhoons. This appeared to be driven by the high growth potential of *L. minor*, since all mesocosm communities with highest realised response imbalance comprised *L. minor* and two other species in combination (Figure SFF). In this way, realised response imbalance provided more explanatory power for stability than either the balance of species responses or the relative abundances of species alone.

We measured population and community resistance to typhoon disturbance as the instantaneous log response ratio between the data collected immediately prior to and following typhoon landfall (Hillebrand et al. 2018). The duration of our experiment did not allow us to consider the longer term recovery trajectories of these communities. However, resistance and recovery often trade off, with resistant communities having slower recovery and *vice versa* (Miller & Chesson 2009; Patrick et al. 2022; but see Poorter et al. 2021). Further, ecological responses are not instantaneous, resulting in transient dynamics and slow recovery in many systems (Hastings et al. 2018), including in response to typhoons (Peierls et al. 2003; Tsai et al. 2011). This has led to calls to integrate longer-term ecological responses to typhoon disturbance (Lin et al. 2020; Stockwell et al. 2020). Our results also support previous work showing species-specific responses to typhoon disturbance (*e.g.*, Ross et al. 2024), in this case likely driven by species’ NO_3_^−^ responses (McCann 2016). The extent to which the structure of species’ environmental responses affects dimensions of stability beyond resistance and temporal stability remains an open question.

We developed a novel machine learning-assisted image classification workflow for estimating the proportional cover of four floating macrophyte species (*Azolla cristata*, *Lemna minor*, *Salvinia molesta*, and *Spirodela polyrhiza*). This workflow, which we call *LeafMosaic* (Mihai 2025), is robust to noise in the form of camera angle, light conditions from weather, blur, and phenotypic plasticity, and can distinguish between two morphologically similar duckweeds, *L. minor* and *S. polyrhiza*. Image classification tasks for floating macrophytes range from measurement of individual fronds for growth in laboratory conditions (Cox et al. 2022; Kose et al. 2023) to broad-scale biomass estimation from satellite images (Castillo et al. 2008; Villa et al. 2018). To our knowledge, *LeafMosaic* is the first workflow to focus on multispecies classification from images of outdoor mesocosm communities, and as such is generalisable to images taken in a range of field conditions. Accordingly, our workflow enhances the tractability of using multispecies competitive communities of floating aquatic macrophytes to study a range of questions in community ecology, including field and semi-controlled experimental studies of competition and coexistence (Armitage & Jones 2019, 2020), diversity-productivity relationships (Couture et al. 2025), response diversity (McCann 2016), and multiple facets of ecological stability (Scheffer et al. 2003; Wang et al. 2025). We successfully applied the *LeafMosaic* workflow to a simple experimental test of the relationship between response diversity and stability, indicating that future studies in this model system hold significant promise.

In sum, we tested whether the balance of floating macrophyte species responses to NO_3_^−^ enrichment predicts aggregate community temporal variability, using experimental mesocosm communities. The balance of species responses was unimportant for community stability, which instead was best explained by species asynchrony and mean population stability. In contrast, when measuring community-level resistance to an unexpected pulse disturbance, a large typhoon, we found that the structure of species responses in the community predicted resistance, but not in the direction we expected. Instead of a balance of positive and negative NO_3_^−^ responses conferring stability, imbalanced responses best predicted community resistance, seemingly driven by positive post-typhoon growth rates of the duckweed *Lemna minor*. Overall our results underscore the importance of the structure of species’ environmental responses in driving stability, as well as the need to consider multiple dimensions of stability in response diversity frameworks.

## Data, Materials, and Software Availability

All data and code needed to reproduce the results of this paper will be archived with the Zenodo repository (proving a DOI to a Github repository). The LeafMosaic machine learning workflow presented here is archived and open source on Zenodo (Mihai 2025).

## Acknowledgements

We thank Aurika Musiienko, Alex Alonso, Damien Morell, and Katie Saunders for help with fieldwork and lab work, Masako Ogasawara for fieldwork permitting, and Sofia van Moorsel for early discussion. This study was supported by a BES Large Grant (LRB22/1007) and a JSPS KAKENHI Grant-in-Aid for Research Start-up 23/22K21332 to SRPJR. All authors were supported by subsidy funding to the Okinawa Institute of Science and Technology Graduate University (OIST). This paper in part results from the activities and support of the Response Diversity Network (https://responsediversitynetwork.githubwebsite/).

## Author contributions

*Samuel R.P-J. Ross*: Conceptualisation; Data curation; Formal analysis; Funding acquisition; Investigation; Methodology; Project administration; Resources; Validation; Visualisation; Writing – original draft. *Alexandru Mihai*: Data curation; Formal analysis; Methodology; Software; Validation; Visualisation; Writing – review and editing. *Chii Kojima*: Data curation; Methodology; Writing – review and editing. *David W. Armitage*: Conceptualisation; Methodology; Resources; Supervision; Validation; Writing – review and editing.

