## Supplementary material for "Species responses to nutrient loading promote resistance but not temporal stability in floating macrophyte communities": _

### Supporting information: Species responses to nitrate addition promote resistance but not temporal stability in floating macrophyte communities

Samuel R.P-J. Ross*, Alexandru Mihai, Chii Kojima, David W. Armitage

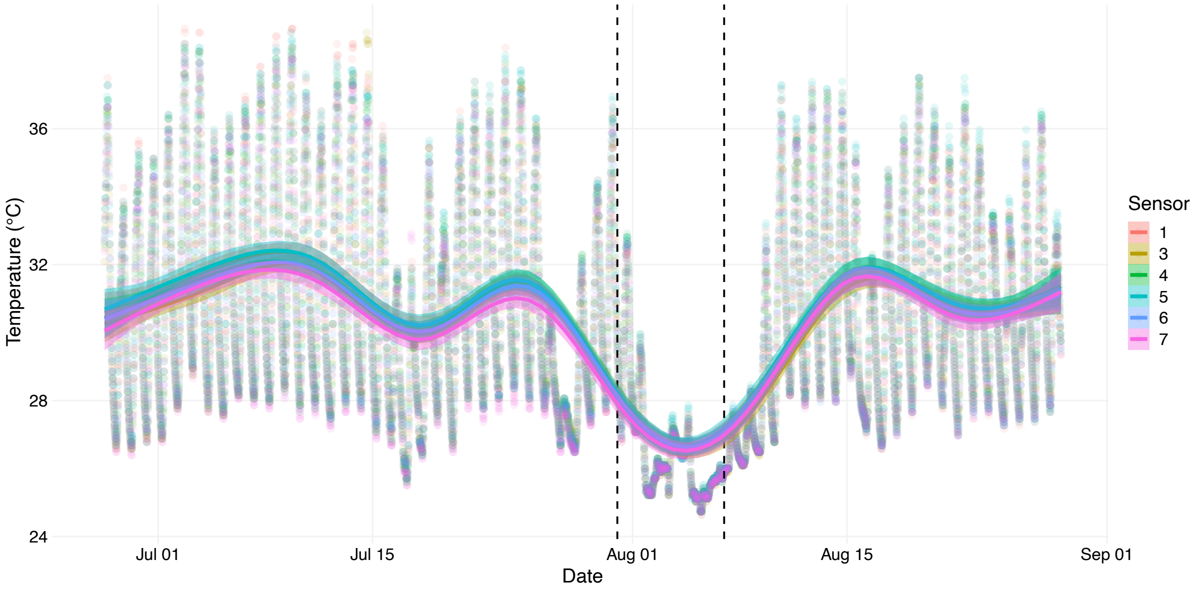


**Figure S1**. Water temperature time series during our experimental period. Water temperature was recorded every 30 minutes in 7 mesocosms using HOBO Pendant Temperature data loggers (model UA-002-64; HOBO, Onset Computer Corporation, Bourne, MA, USA). Temperatures did not differ significantly between mesocosms, and ranged from 24.6 – 38.9 °C. Landfall of typhoon Khanun around 31 Jul – 6 Aug (denoted by dashed lines; JMA 2023) reduced water temperatures and daily temperature variability.

**Table S1**. Nutrient additions to each mesocosm, including details of nitrate treatments and scaled amount of chemicals added to all 69 buckets combined. H-40 growth medium and Iron & micronutrients were added at the start of the experiment and again after 4 and 8 weeks. $NO_3^-$ treatments were maintained by weekly additions of crystalline $NaNO_3$ (see **Methods**).

| H-40 growth medium | g/bucket | mg/bucket | Total amount (g/69 buckets) |
|---|---|---|---|
| $CaCl_2.2H_2O$ | 9.86 | 9,856 | 680.1 |
| $MgSO_4.7H_20$ | 7.76 | 7,762 | 535.6 |
| NaCl | 0.16 | 157 | 10.82 |
| $KH_2PO_4$ | 0.46 | 460.8 | 31.8 |
| $K_2HPO_4.3H_2O$ | 0.024 | 24.2 | 1.66 |
| KCl | 4.44 | 4,440.2 | 306.4 |

| Iron & micronutrients | L/bucket | mL/bucket | Total amount (mL/69 buckets) |
|---|---|---|---|
| フルボ鉄 | 0.0002 | 0.2 | 13.8 |

| $NO_3^-$ Treatments | g/bucket | mg/bucket | Total amount (g/23 buckets) |
|---|---|---|---|
| $NaNO_3$: Low | 0.025 | 24.95 | 0.575 |
| $NaNO_3$: Medium | 0.09 | 89.94 | 2.07 |
| $NaNO_3$: High | 0.2 | 200 | 4.6 |

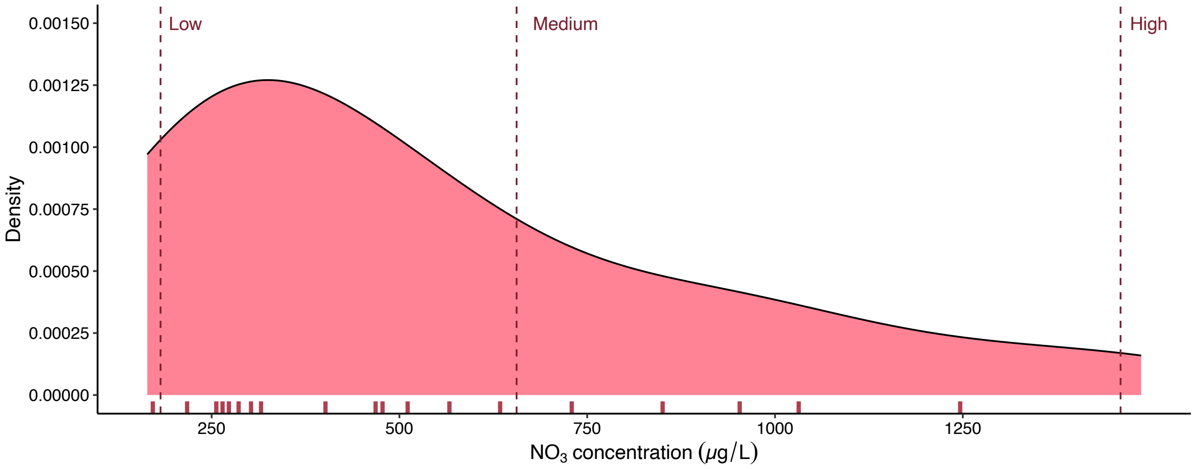


Figure S2. Distribution of $NO_3^-$ concentration values for water collected from agricultural ponds and ditches around Okinawa (see main text). Densities represent analysed water samples, and dashed vertical lines indicate our $NO_3^-$ concentration treatment levels used in the mesocosm experiment. X-axis shows the 95th percentiles at 5% intervals from 5% to 95%. Our low $NO_3^-$ concentration treatment falls around the 5th percentile, the medium treatment around the 70th percentile, and the high treatment beyond the 95th percentile. Accordingly, the high and low treatments represent extreme (but realistic) $NO_3^-$ scenarios, while the medium treatment is closer to average $NO_3^-$ concentrations in agricultural ditches and ponds around Okinawa.

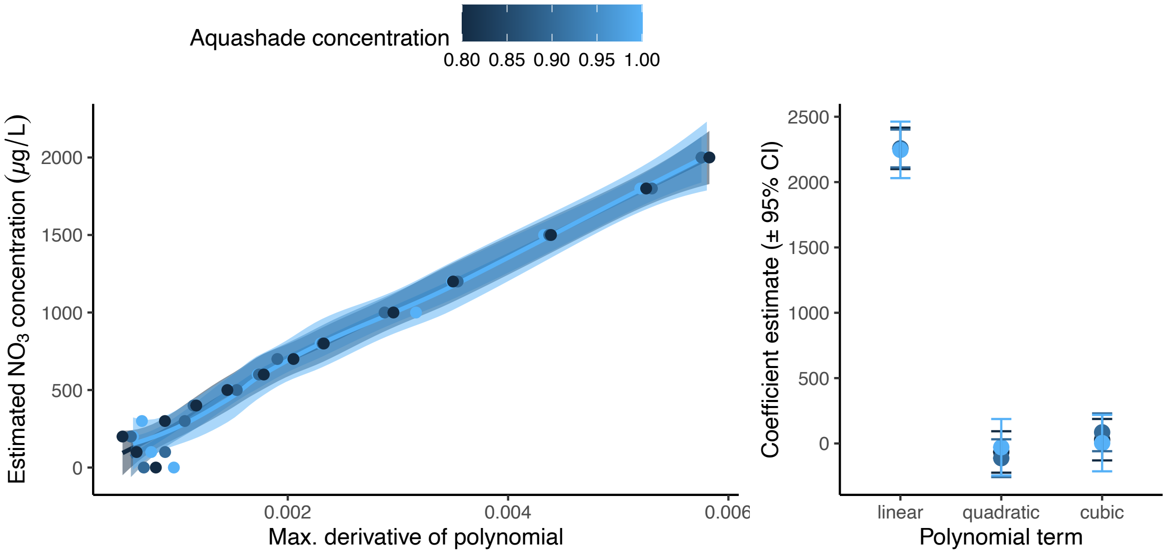


Figure S3. Effects of Aquashade concentration on of $NO_3^-$ concentration in laboratory assay samples. We used the second derivative calculation method for direct spectrophotometric analysis of aqueous nitrates (Olsen 2008) to estimate $NO_3^-$ concentration along a gradient of known $NO_3^-$ concentration (0–2000 μg $L^{-1}$) at relative Aquashade concentrations (0.8–1.00, v/v, where 1.00 denotes undiluted stock solution). Left panel shows the relationship between the maximum derivative of the fitted polynomial function across this assay. Polynomial response functions were best described by a linear term (right panel) and were fitted separately for each Aquashade concentration. Shaded regions are 95% confidence intervals around fitted relationships. Right panel shows estimated coefficients (± 95% confidence intervals) for linear, quadratic, and cubic polynomial terms included in fitted models. The relationship was best explained by the linear term, and in no cases were differences between Aquashade detected based on non-overlapping 95% confidence intervals. Variation in Aquashade concentration therefore did not significantly affect estimated $NO_3^-$ concentrations across the tested range.

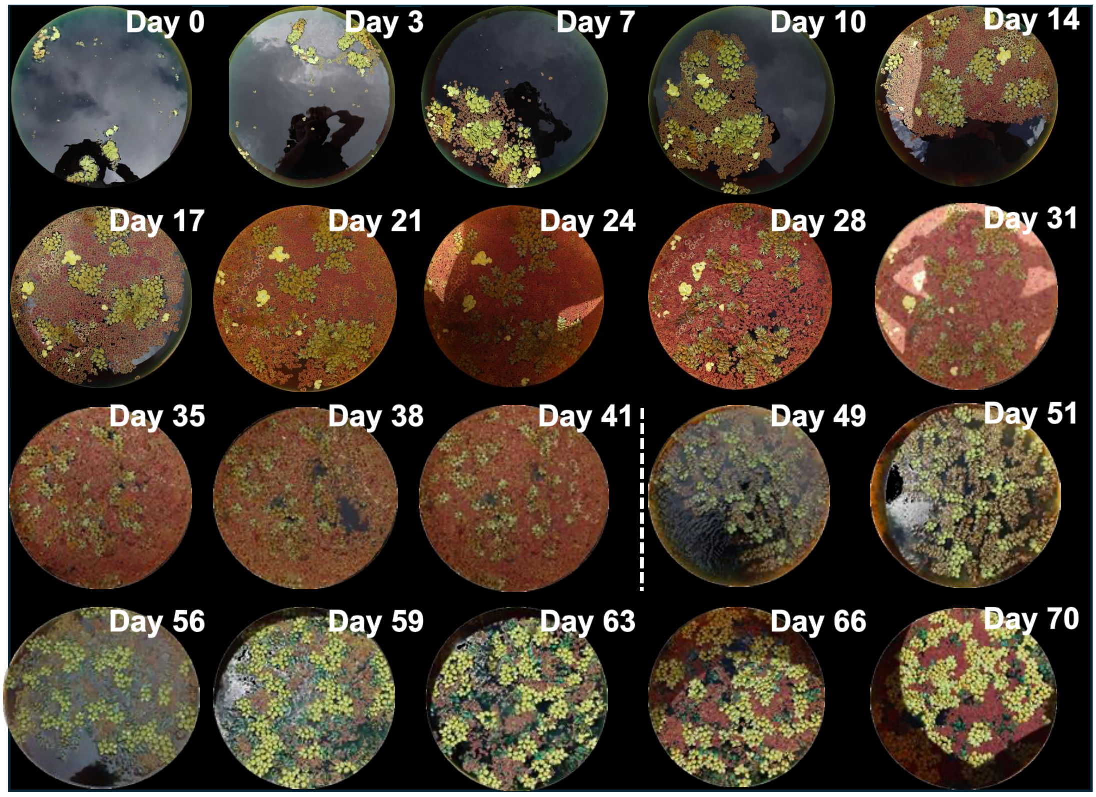


Figure S4. Photographic time series illustrating community compositional change in mesocosm #1 across the 70-day experimental period. The passing of a large typhoon is marked with a dashed line. Note that both *Azolla cristata* and *Salvinia molesta* exhibited phenotypic change during the experiment; *A. cristata* became redder in some mesocosms, and *S. molesta* varied in shape and colour.

Table S2. No meaningful effect of shifting tile location on macrophyte classification results. We shifted the starting location of our 100-by-100-pixel tiles by 25 and 50 pixels in their x- and y- coordinates. We found minimal effect on model classification results from DenseNet (measured here before downstream corrections such as time series smoothing; see Table S7). Data are species-specific proportional cover from image tile classifications. Species typically varied less than 1 percentage point, and total macrophyte cover differed by less than 2 percentage points in all cases.

| Shift (pixels) | *A. cristata* cover | *L. minor* cover | *S. molesta* cover | *S. polyrhiza* cover | *Total* |
|---|---|---|---|---|---|
| None | 0.076 | 0.276 | 0.475 | 0.015 | 0.842 |
| 25 X | 0.086 (+0.01) | 0.256 (-0.02) | 0.482 (+0.007) | 0.009 (-0.006) | 0.833 |
| 25 Y | 0.066 (-0.01) | 0.288 (+0.012) | 0.468 (-0.007) | 0.016 (+0.001) | 0.838 |
| 50 X | 0.073 (-0.003) | 0.265 (-0.011) | 0.482 (+0.007) | 0.009 (-0.006) | 0.829 |
| 50 Y | 0.073 (-0.003) | 0.265 (-0.011) | 0.475 (0) | 0.012 (-0.003) | 0.825 |

#### Supporting Methods

***Training dataset composition***

Unsupervised clustering (UMAP + HDBSCAN) applied to CNN-extracted features from 40 representative photographs produced visually coherent groups of image tiles, which were then manually verified and assigned to species-specific classes (see **Methods**). Given the pronounced phenotypic variation in *Salvinia molesta* (green vs. brown leaves) and *Azolla cristata* (green vs. red fronds), each of these species was split into two phenotype sub-classes for training (Table S3). Together with Lemna minor, Spirodela polyrhiza, and two water classes (dark water and light water/glare), this yielded eight classification classes.

Table S3. Composition of the verified training dataset. Phenotype sub-classes are shown for *S. molesta* and *A. cristata*. Tiles were retained only if a single species occupied >50% of the tile area and the assigned phenotype was visually consistent across clusters.

| Species/Class | Phenotype | Tiles |
|---|---|---|
| *Azolla cristata* | Green | 3,794 |
| | Red | 3,599 |
| | Subtotal | 7,393 |
| *Lemna minor* | — | 2,854 |
| *Salvinia molesta* | Dark (brown) | 2,364 |
| | Light (green) | 2,510 |
| | Subtotal | 4,874 |
| *Spirodela polyrhiza* | — | 2,560 |
| Water (background) | — | 2,073 |
| **Total** | | **19,754** |

##### *Model architecture comparison and accuracy*

We trained 15 separate classification models spanning all pairwise-to-full combinations of our four target species, using two convolutional neural network backbones: DenseNet121 and VGG16, both initialised with ImageNet weights and fine-tuned classifier heads. Models were evaluated on a held-out validation set comprising 20% of the data. The primary performance metric was categorical accuracy on the validation set ($acc_{val}$), defined as the fraction of validation tiles assigned the correct species label across all classes (Tables S4 and S5). To select the best training checkpoint and compare architectures, we computed a ranking score for each epoch combining validation accuracy, a normalised validation loss term, and a one-sided overfitting penalty:

$$score = 0.75 \cdot acc_{val} + 0.10 \cdot L_{val} - 0.15 \cdot max(0, L_{val} - L_{train})$$

where $L_{val}$ is the min–max normalised validation loss:

$$L_{val} = 1 - \frac{L_{val} - L_{min}}{L_{max} - L_{min}}$$

inverted so that lower loss yields a higher contribution, and max(0, $L_{val}$ – $L_{train}$) penalises epochs where validation loss exceeds training loss, indicating overfitting. The weighting scheme prioritises validation accuracy as the principal criterion (0.75), treats loss as a secondary quality signal (0.10), and applies a moderate overfitting penalty (0.15). The epoch maximising this score was selected as the best checkpoint for each model.

Within the training-testing framework, validation accuracy was highest for single-species classifiers (mean 93.3%) and declined modestly with increasing species richness, reaching 90.0% for four-species mixtures. This decline is expected as visual confusion between co-occurring species increases with community complexity, particularly between the visually similar duckweeds *L. minor* and *S. polyrhiza*, and between *S. molesta* and *A. cristata*, which can have similar brown or green phenotypes under different nutrient conditions. Nine of the 15 trained models exceeded 95% validation accuracy, and no model yielded below 85% accuracy.

Table S4. Mean classification accuracy (%) by architecture, averaged across all 15 species-combination models.

| Architecture | Mean accuracy (%) | Notes |
|---|---|---|
| DenseNet121 | 92.0 | Most versatile across species combinations |
| VGG16 | 90.7 | Competitive but higher overfitting tendency |
| **Overall mean** | **91.4** | **Across all model-species combinations** |

Table S5. Per-species classification accuracy (%) averaged across all architectures and species-combination models in which each species appeared. Standard deviation reflects variation across models of different species richness levels.

| Species | Accuracy % (mean ± SD) |
|---|---|
| *Lemna minor* | 93.0 ± 2.8 |
| *Spirodela polyrhiza* | 92.1 ± 3.2 |
| *Azolla cristata* | 91.4 ± 3.5 |
| *Salvinia molesta* | 89.1 ± 4.1 |

##### *Within-tile macrophyte cover calculation and post-processing*

For each classified tile, we estimated the proportion of the tile area occupied by macrophyte vegetation using colour-space filtering in HSV and LAB. This within-tile cover estimate was used to filter out tiles with < 0.5 proportional plant cover, which were excluded from species classification.

The within-tile cover mask was constructed by combining three hue-based masks with a lightness constraint (Table S6). Each tile was converted from BGR to HSV and LAB colour spaces. Pixels falling within any of the three hue ranges and exceeding the minimum lightness threshold were classified as plant material. The final macrophyte mask M was computed as:

$$M = (M_{green} \cup M_{yellow} \cup M_{brown}) \cap M_{lightness}$$

Table S6. HSV and LAB color-space parameters used to estimate within-tile macrophyte cover per tile. H, S, and V denote hue, saturation, and value channels of the HSV color space; L denotes the lightness channel of the LAB color space. The three hue masks capture the dominant colour ranges of our four macrophyte species across their observed phenotypes.

| Mask | H range | S range | V range | L range (LAB) |
|---|---|---|---|---|
| Lightness gate | — | — | — | 35–230 |
| Green | 30–85 | 60–255 | 60–255 | (above) |
| Yellow | 18–35 | 80–255 | 100–255 | (above) |
| Brown | 5–18 | 100–255 | 50–180 | (above) |

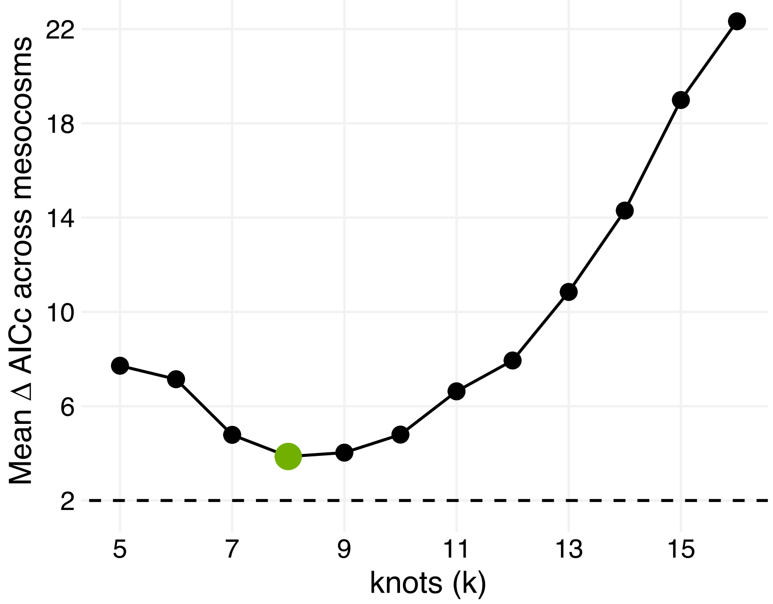


Figure S5. Guassian process smooth performance across mesocosms. Model selection results for basis functions (*k*) in a generalised additive model with beta family and logit link function. The model fitted a gaussian process smoother with different numbers of basis functions. Optimal *k* (8) was defined based on lowest ΔAICc across all mesocosms on average. The final choice of *k* had, on average across all mesocosms, ΔAICc = 3.87. None of the *k* values tested fell within the range of ΔAICc ≤ 2 averaged across all mesocosms.

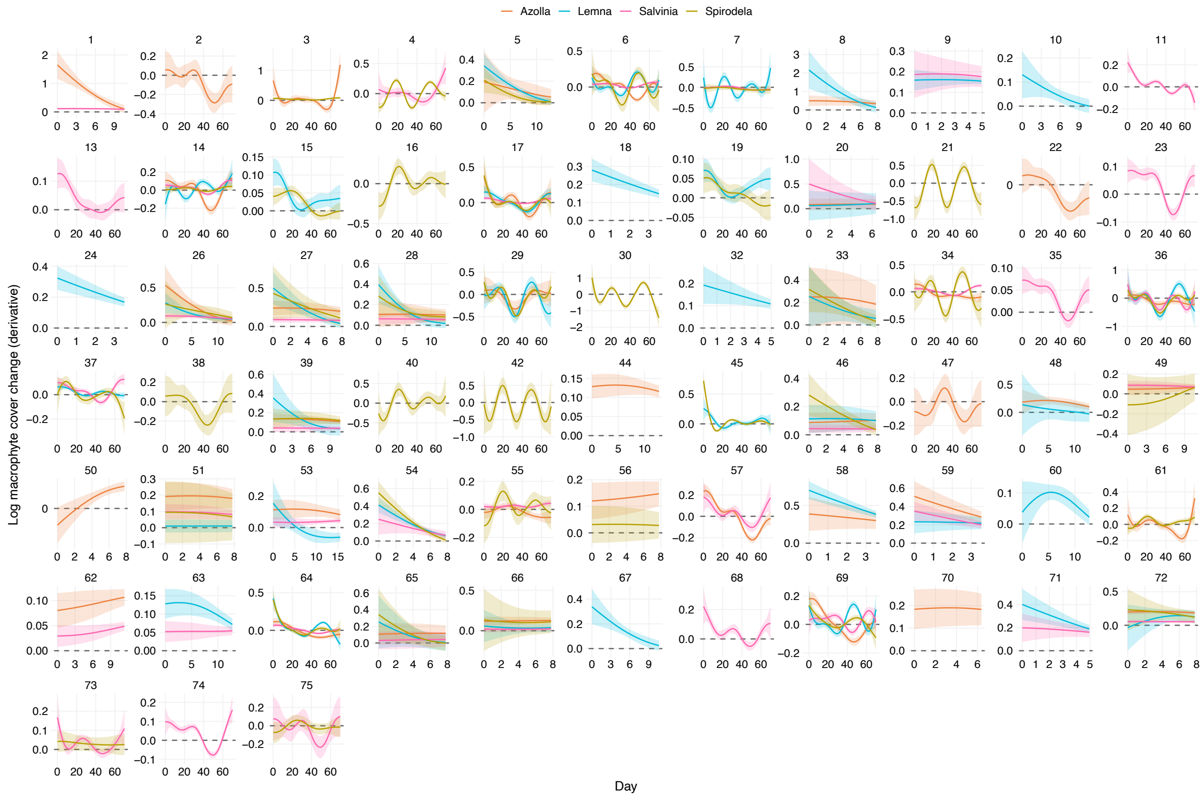


Figure S6. Changes in log macrophyte cover for growth rate calculation for each mesocosm. We modelled time (day since start) as a Gaussian process smooth with $k$ = 8 (Figure S5), and here show the first derivatives of these smooths with 95% confidence intervals. Positive values (above the dashed line) indicate an increase in the modelled relationship between macrophyte cover and time, negative values indicate a decrease. We took growth rates as the arithmetic mean of these derivative time series. Note that the number of days on the x-axis varies as time series were cut to only include dates before total macrophyte cover exceeded 90% of the water surface. Colours represent different species: *Azolla cristata* in orange, *Lemna minor* in blue, *Salvinia molesta* in pink, and *Spirodela polyrhiza* in yellow.

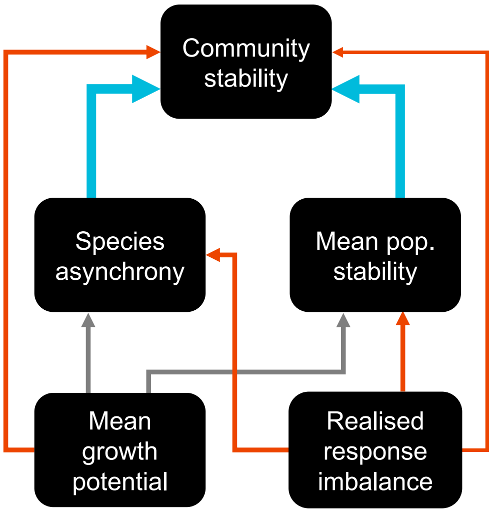


Figure S7. Hypothesised relationships in SEM analysis. Red arrows represent hypothesised negative relationships, blue arrow hypothesised positive relationships, and grey arrows paths for which we had no strong hypotheses but which were included in the model based on posterior checks. Arrow size scales with the hypothesised strength of predicted relationships. We expected that communities with faster growing species on average, should be less stable, though we had no specific hypothesis regarding the relationship between mean growth potential and species asynchrony or mean population stability (these SEM paths were added following significant tests of directed separation). We expected species asynchrony and mean population stability to both have a positive effect on community stability (Thibaut & Connolly 2013; Sasaki et al. 2019). We expected response diversity—here measured as the realised response imbalance (Polazzo et al. 2025)—to be stabilising (Mori et al. 2013; Ross et al. 2023). Here higher realised response imbalance represents communities with more skewed $NO_3^-$ concentration responses, and therefore is not directly equivalent with response *diversity*; we expect a balance of positive and negative responses to be stabilising (Yachi & Loreau 1999; Ross et al. 2023; Polazzo et al. 2025), so we hypothesised a negative link between realised response imbalance and community stability. Communities with more balanced $NO_3^-$ concentration responses (low realised response imbalance) are expected to arise from either of two mechanisms: high species asynchrony or high mean population stability (Ross & Sasaki 2024; Polazzo et al. 2025), so we anticipated negative links between realised response imbalance and each of these mechanisms. In this way, we expected that response imbalance more strongly acts as an indirect stability mechanism through its direct effects on species asynchrony and mean population stability, rather than a strong direct effect of response imbalance (Sasaki et al. 2019; White et al. 2023; Ross & Sasaki 2024; Polazzo et al. 2025).

Table S7. Model architectures and post-processing filters for validation against manually classified image tile counts. Species names indicate cases for which the model architecture or post-processing adjustment performed best for that species based on model slope, intercept, and variance explained (see **Methods**). Confusion adjustment only applies to cases where *S. polyrhiza* and *L. minor* are confused. See Figure S8 for performance of final chosen models.

| Model | Time series smooth | Within-tile cover | Confusion adjustment |
|---|---|---|---|

| | | | |
|---|---|---|---|
| VGG | None | None<br>*L. minor*<br>*S. polyrhiza* | None |
| DenseNet<br>*L. minor*<br>*S. polyrhiza* | Loess smooth ($\alpha$ = 0.3)<br>*A. cristata*<br>*L. minor*<br>*S. molesta*<br>*S. polyrhiza* | <0.3 cover removed<br>*A. cristata*<br>*S. molesta* | <0.05 difference removed<br>*S. polyrhiza* |
| Model average<br>*A. cristata*<br>*S. molesta* | | <0.5 cover removed | <0.05 difference reassigned<br>*L. minor*<br><br><0.1 difference removed<br><0.1 difference reassigned |

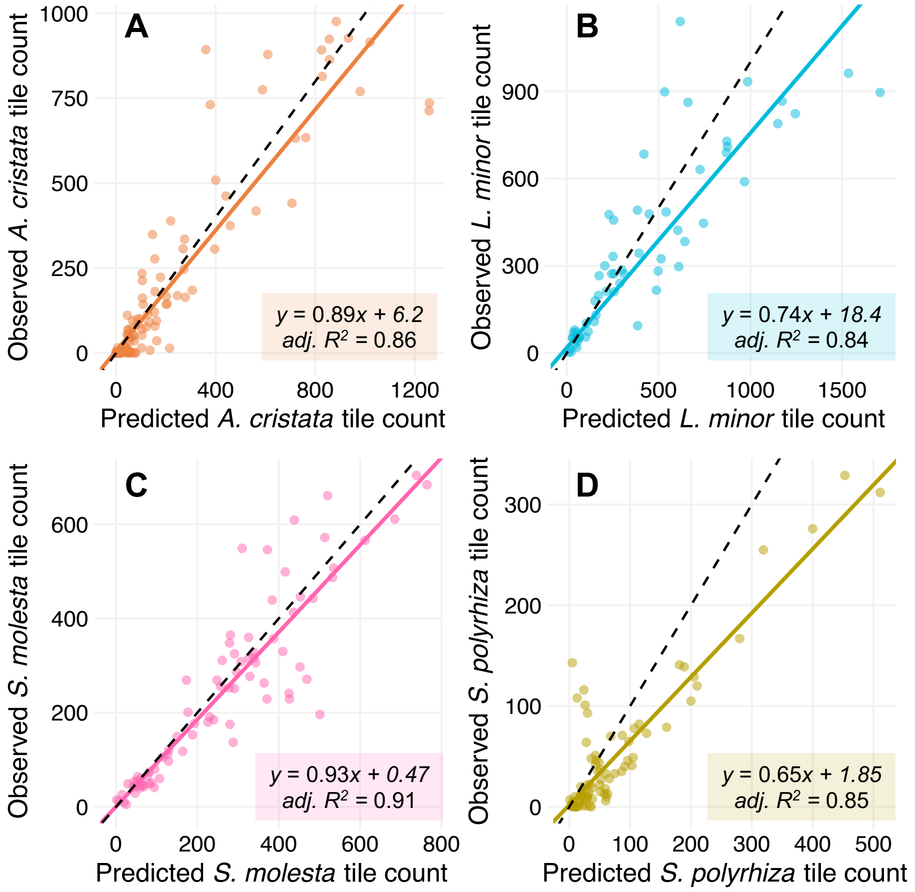


**Figure S8**. Final machine learning classifier validation plots. Plots show predicted image tile counts from the best model for each species (Table S7) across 160 photographs from 8 mesocosms (a validation set, see **Methods**) on x-axes and the manually observed tile counts on Y-axes for each of (A) *Azolla cristata*, (B) *Lemna minor*, (C) *Salvinia molesta*, and (D) *Spirodela polyrhiza*. Model performance was assessed via fit of each line and ability to reproduce temporal dynamics (Figure S9).

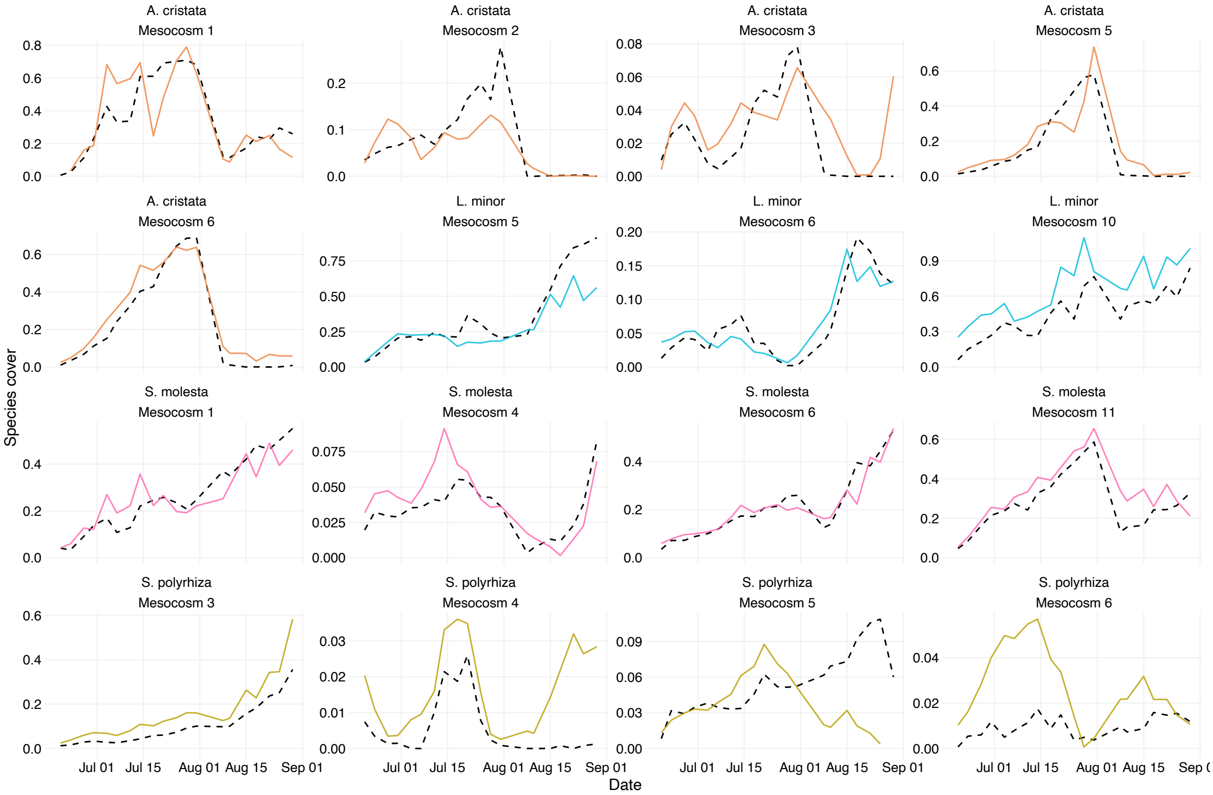


Figure S9. Final machine learning classifier validation time series. Plots show predicted image tile counts (coloured lines) from the best model for each species (Table S7) versus manually observed tile counts (dashed lines) through time for different species-by-mesocosm combinations from the validation set (see **Methods**). Note that y axis values differ substantially, such that mismatches were never large (e.g. for *S. polyrhiza*). Model performance was assessed via fit of each line (Figure S8) and ability to reproduce temporal dynamics.

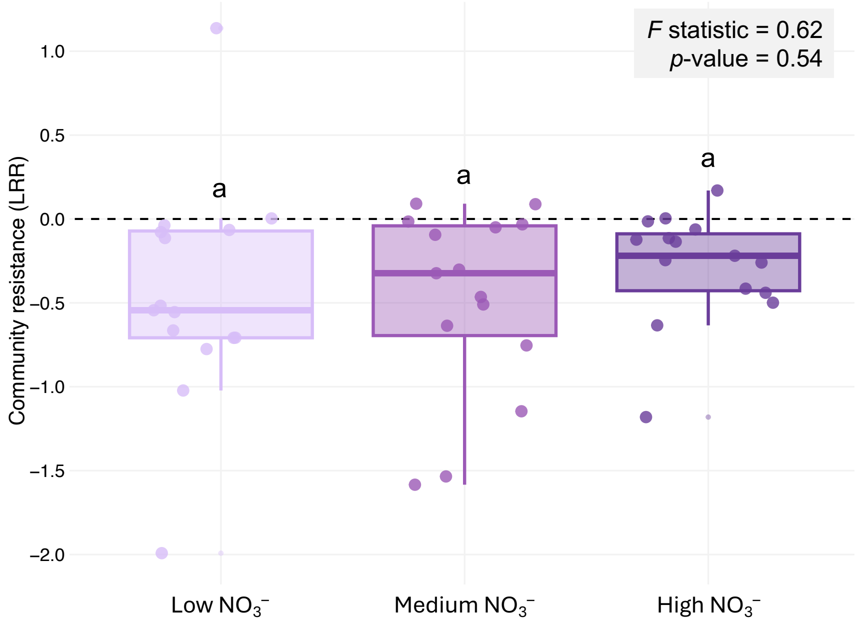


Figure S10. Nonsignificant effect of nitrate enrichment treatment on community resistance to discrete typhoon disturbance, measured as the log response ratio of total macrophyte cover before versus after typhoon disturbance. Values above zero indicate a post-typhoon increase in total macrophyte cover, while values below zero indicate a decrease. Multivariate analysis of variance (ANOVA) revealed no difference between $NO_3^-$ treatments in their community resistance ($ANOVA_{2,42}$: $F = 0.62$, $p = 0.54$).

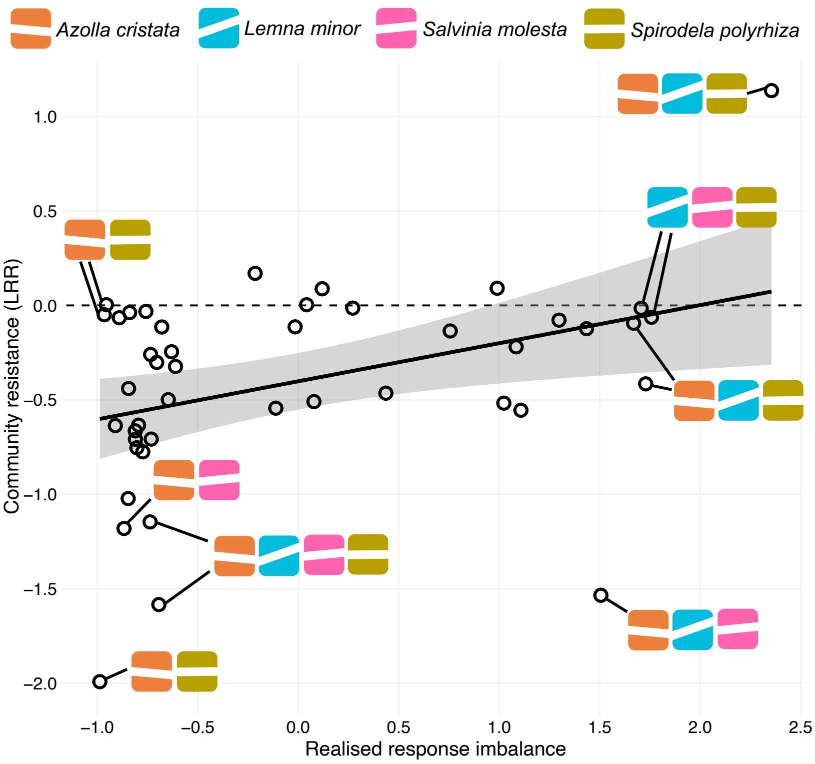


Figure S11. Figure illustrating the significant positive relationship between realised response imbalance (Polazzo et al. 2025) and community resistance to typhoon disturbance (Figure 5A). Key points for understanding the relationship are annotated with species composition. White lines through coloured boxes represent the corresponding species' growth response to nitrate ($NO_3^-$) enrichment from which response imbalance was calculated (see Figure 3), to visually illustrate the response diversity of each mesocosm community. Note that values of response imbalance were scaled by the relative biomass (macrophyte cover) contribution of each species, making them *realised* response imbalance values. We find that highest response imbalance results from communities where the species with more extreme $NO_3^-$ responses (*Lemna minor* and to a lesser extent *Salvinia molesta*) comprised a greater proportion of total macrophyte cover through time, meaning these stronger responding species represented the dominant response within a given mesocosm community.

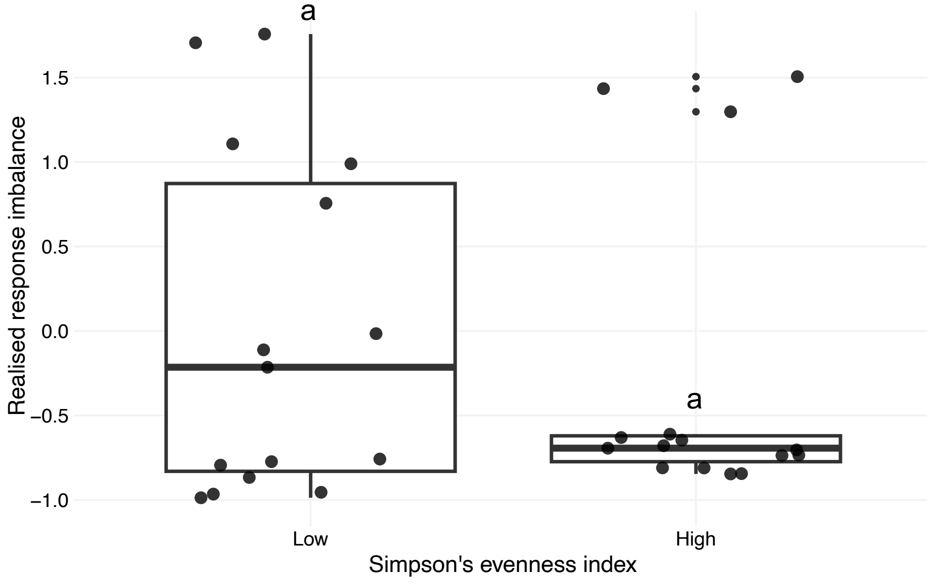


Figure S12. No significant difference in realised response imbalance between high versus low evenness communities, measured as Simpson diversity. Low and high categories were split as the lower and upper 33% of the range of Simpson evenness values, respectively. A T-test revealed no significant difference (T-test: $t$ = 0.84, $df$ = 27.6, $p$ = 0.41).

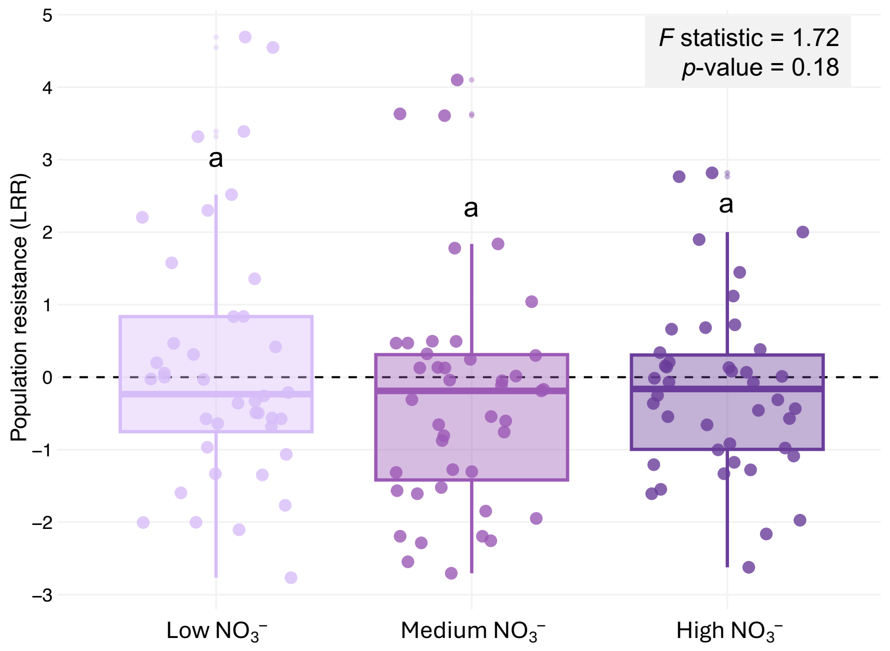


Figure S13. Nonsignificant effect of nitrate enrichment treatment on population resistance to discrete typhoon disturbance, measured as the log response ratio of macrophyte cover per species (aggregated here) before versus after typhoon disturbance. Values above zero indicate a post-typhoon increase in total macrophyte cover, while values below zero indicate a decrease. Multivariate analysis of variance (ANOVA) revealed no difference between $NO_3^-$ treatments in their population resistance when ignoring species effects ($ANOVA_{2,113}$: $F$ = 1.72, $p$ = 0.18).
